# Novel in-silico predicted matrikines are differential mediators of in vitro and in vivo cellular metabolism

**DOI:** 10.1101/2023.03.17.533127

**Authors:** Nathan Jariwala, Matiss Ozols, Alexander Eckersley, Bezaleel Mambwe, Rachel E B Watson, Leo Zeef, Andrew Gilmore, Laurent Debelle, Mike Bell, Eleanor J Bradley, Yegor Doush, Carole Courage, Richard Leroux, Olivier Peschard, Philippe Mondon, Caroline Ringenbach, Laure Bernard, Aurelien Pitois, Michael J Sherratt

**Affiliations:** Division of Cell Matrix Biology & Regenerative Medicine, School of Biological Science, Faculty of Biology, Medicine and Health, The University of Manchester, Manchester, UK; Department of Human Genetics, Wellcome Sanger Institute, Genome Campus, Hinxton, UK; British Heart Foundation Centre of Research Excellence, University of Cambridge, Cambridge, UK; Division of Musculoskeletal & Dermatological Sciences, Faculty of Biology, Medicine and Health, The University of Manchester, Manchester, UK; Bioinformatics Core Facility, Faculty of Biology, Medicine and Health, The University of Manchester, Manchester, UK University of Manchester, UK; Wellcome Centre for Cell Matrix Research, Division of Cancer Sciences, Faculty of Biology, Medicine and Health, University of Manchester, UK; UMR CNRS 7369 MEDyC, Université de Reims Champagne Ardenne, UFR Sciences Exactes et Naturelles, SFR CAP Santé, Moulin de la Housse, 51687 Reims cedex 2, France; No7 Beauty Company, Walgreens Boots Alliance, Nottingham, UK; Sederma, 29 rue du Chemin Vert, Le Perray en Yvelines, 78612, France

**Author notes:** These authors contributed equally: Nathan Jariwala, Matiss Ozols.

**Keywords:** extracellular matrix, peptides, matrikines, skin, ageing, proteases, therapeutic discovery

## Abstract

The exogenous application of small peptides can beneficially affect clinical skin appearance (wrinkles) and architecture (collagen and elastic fibre deposition and epidermal thickness). However, the discovery of new bioactive peptides has not been underpinned by any guiding hypothesis. As endogenous extracellular matrix (ECM)-derived peptides produced during tissue remodelling can act as molecular signals influencing cell metabolism, we hypothesised that protease cleavage site prediction could identify putative novel matrikines with beneficial activities. Here, we present an *in silico* to *in vivo* discovery pipeline, which enables the prediction and characterisation of peptide matrikines which differentially influence cellular metabolism *in vitro*. We use this pipeline to further characterise a combination of two novel ECM peptide mimics (GPKG and LSVD) which act *in vitro* to enhance the transcription of ECM organisation and cell proliferation genes and *in vivo* to promote epithelial and dermal remodelling. This pipeline approach can both identify new matrikines and provide insights into the mechanisms underpinning tissue homeostasis and repair.

## Introduction

Extracellular matrices (ECMs) of mammalian tissues play important roles in mediating tissue structure, mechanical properties and cellular phenotype (Theocharis et al., 2016). However, aberrant and progressive remodelling of ECM-rich tissues is a key feature in the pathology and ageing of many organs, including articular cartilage (Bolduc et al., 2019), arteries (Lacolley et al., 2018) and skin (Wilkinson and Hardman, 2021). Whilst some ECM proteins, such as newly synthesised collagen I in tendon, are maintained through a circadian cycle (Chang et al., 2020), the longevity of other ECM components makes them vulnerable to oxidative damage (Sander et al., 2002), pathological cross-linking (Moldogazieva et al., 2019) and protease-mediated degradation (Freitas-Rodriguez et al., 2017). In particular, many structural ECM components such as elastin (Shapiro et al., 1991), aggrecan and collagen (Sivan et al., 2006) persist in tissues for decades. The relative susceptibility of individual ECM proteins to degradation may be determined not only by their longevity but also abundance, tissue location (for example in the ultraviolet radiation [UVR]-exposed papillary dermis of photoaged skin (Watson et al., 1999)) and biochemistry (where amino acid composition mediates susceptibility to UVR and oxidation (Hibbert et al., 2015)). Crucially, ECM degradation may not only impair function but also release peptide fragments, known as matrikines, or reveal previously shielded active sites known as matricryptins (Davis et al., 2000) with cell signalling capabilities (Maquart et al., 1999).

Early research into the bioactivity of ECM fragments focussed on elastin-derived peptides, which exhibit diverse actions against stromal, endothelial and immune cells (Duca et al., 2004), but it is clear that matrikines can be liberated from multiple ECM proteins. For example, the collagen IV NC1 domain fragments canstatin (Kamphaus et al., 2000) and arresten (Nyberg et al., 2008) are anti-angiogenic, tumour suppressive and able to regulate apoptosis. Furthermore, a smaller peptide fragment of canstatin (amino acids 78-86; known as Cans) also demonstrates biological activities, including inhibiting migration and inducing apoptosis in tumour cells (Chamani and Zamani, 2022). The action of matrikines may not always be clinically beneficial, with the collagen-derived matrikine PGP promoting chronic inflammation and pulmonary fibrosis (Bras and Frangogiannis, 2020), although conversely collagen I (C-1158/59 (Lindsey et al., 2015)) and XVIII (endostatin (Isobe et al., 2010)) matrikines have been shown to reduce cardiac fibrosis. These studies, and many others, demonstrate that endogenously generated ECM peptides can act as matrikines, influencing tissue physiology and, ultimately, function. However, they also suggest that the application of exogenous ECM-derived peptides may beneficially affect tissue homeostasis.

The accessibility of human skin, which enables characterisation of ageing (den Dekker et al., 2013) and repair (Watson et al., 2009), its susceptibility to molecular damage (Ozols et al., 2021b) and its cellular and extracellular composition, which is similar to other connective tissue-rich organs, make it an excellent system in which to study the action of exogenous peptides in humans. Human skin may be subject to both intrinsic (passage of time) and extrinsic (action of exogenous factors, often ultraviolet radiation; UVR) ageing (Gu et al., 2020). Whilst there are differences in the clinical manifestations of these ageing processes, both affect the epidermis and dermal ECM proteins (Naylor et al., 2011). In our recent review of bioactive peptides used within skin anti-ageing cosmeceuticals (Jariwala et al., 2022), we identified 35 peptides whose sequences were found in at least one human protein. Many of these peptides were short (di- or tri-peptides), and were usually modified with a palmitoyl chain to aid penetration through the skin barrier (Choi et al., 2014). Comparison of relative activity of these putative ECM matrikines is difficult given the disparity of outcome measures and model systems employed in published studies and, in some cases, the inclusion of the peptide in a complex formulation. However, the beneficial effects of such treatments are clear with, for example, enhanced collagen synthesis *in vitro* (Maquart et al., 1988) and enhanced fibrillin-rich microfibril deposition and wrinkle reduction *in vivo* (Watson et al., 2009) being reported for the peptide palmitoyl-GHK. The short amino acid sequence of this peptide, along with that of other biologically active tri-peptides, means that this peptide, and others which show activity, are commonly found in hundreds or thousands of human proteins (Jariwala et al., 2022). Despite the success of some peptides in inducing clinically discernible benefits in aged skin, to date, there has been no published conceptual framework to guide the prediction and characterisation of new bioactive therapeutic peptides. Instead, new peptide discovery has relied on serendipity, or inferences based on protein active sites and the chemical modification of existing peptides (Leroux et al., 2020).

In this study, we test the hypothesis that small bioactive peptides (matrikines) can be predicted by the *in silico* digestion of dermal proteins via action of ECM proteases. In contrast to enzymes such as trypsin, where cleavage sites can be predicted with a great deal of certainty (Manea et al., 2007), identifying putative cleavage sites of endogenous tissue ECM proteases (such as members of matrix metalloproteinases [MMPs] and cathepsins) requires the use of machine learning algorithms which can predict cleavage sites in protein sequences (Ozols et al., 2021a; Song et al., 2012). We have established a new discovery pipeline in which potential peptide matrikines were predicted and synthesised (following selection based on size, solubility and suitability for manufacturing; Fig. 1a) and screened for *in vitro* cell culture toxicity and biological activity (targeted immunohistochemical markers, and transcriptome and proteome discovery; Fig. 1b). A peptide combination was then progressed to an *in vivo* occluded patch test in human volunteers that simulates the longer-term skin rejuvenation potential of topically applied compounds (Watson et al., 2008) (Fig. 1c). The identification of new matrikines has the potential not only to provide insights into tissue pathology, but also to translate to actives with utility in skin (Cole et al., 2018) and potentially other diverse tissues (such as brain (Reed et al., 2019), muscle (Pavan et al., 2020), heart (Hardy et al., 2019) and liver (Mohammed et al., 2021)) in which treatment of aberrant ECM-remodelling is a pressing and unmet clinical need.

**Figure 1.**
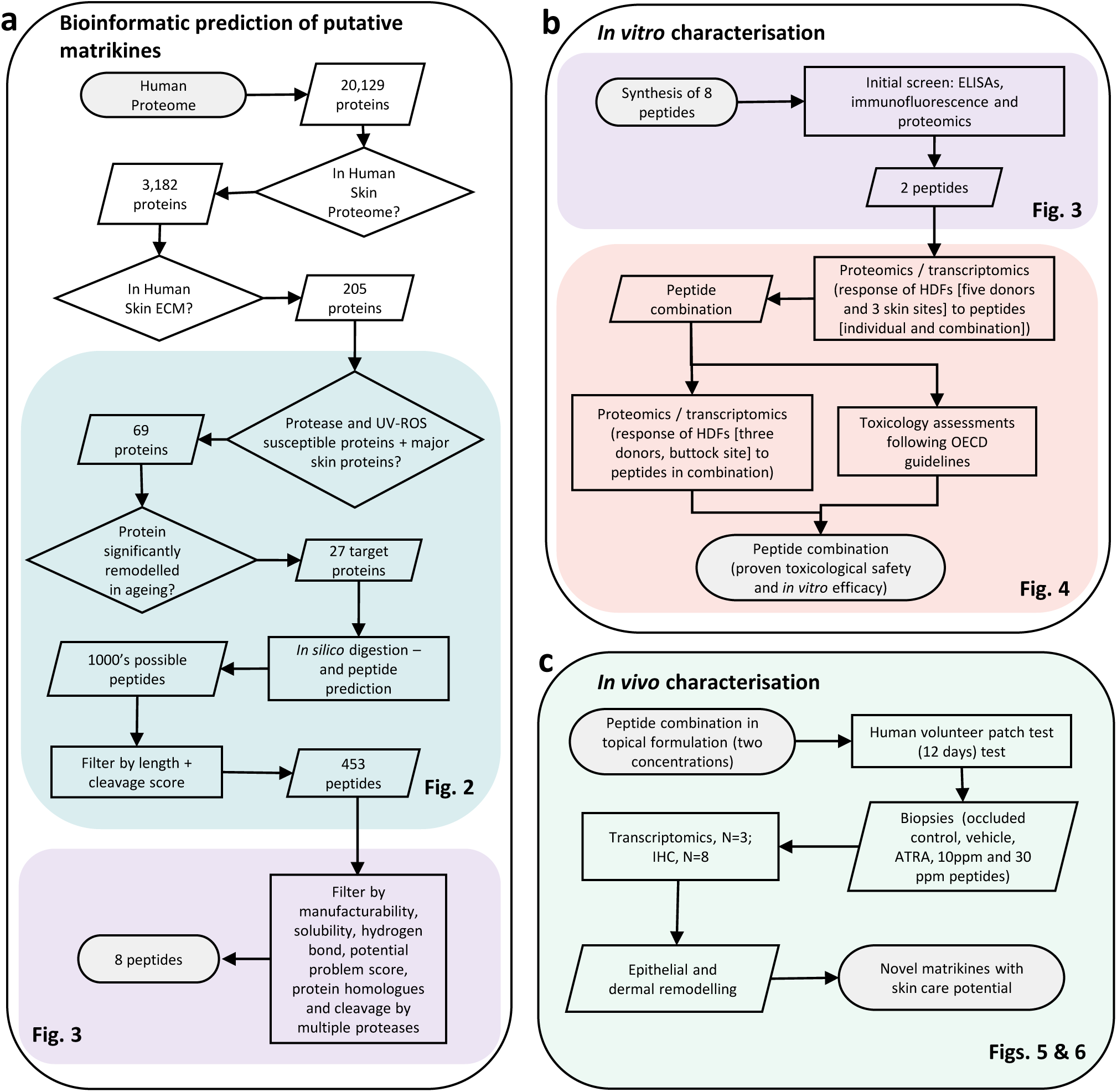
*In silico* to *in vivo* discovery pipeline. (a) Bioinformatic prediction of putative matrikines. The entire human proteome was filtered to identify 205 human skin ECM proteins. An initial protein cohort (key structural ECM and/or higher predicted susceptibility to proteases and UVR/ROS) was further filtered based on reported age-related remodelling to a target cohort of 27 proteins. This cohort was subjected to *in silico* digestion and peptide prediction. The 453 tetra-peptides were then assessed for potential issues with manufacture yielding 8 peptides for synthesis and characterisation. (b) *In vitro* characterisation. The activity of these 8 peptides was assessed by immunological and proteomic screens and 2 peptides were selected for further characterisation. Two rounds of proteomic and transcriptomic assays established that these peptides induced differential and synergistic effects on cell physiology. Toxicological assessment established the safety of the peptide combination in a formulation for topical *in vivo* testing in human skin. (c) Human volunteers were treated with a positive control (ATRA), the vehicle and the peptide formulation at two concentrations (10ppm and 30ppm) for 12 days. Histological, immunohistochemical and transcriptomic analysis established that the peptide combination exerted beneficial effects on key biomarkers of skin photoageing and upregulated expression pathways. Experimental data relating to each step in the discovery pipeline is indicated by colour-coded regions for each figure.

## Results

### Selecting target proteins which are likely to be sources of matrikines *in vivo*

As matrikines are most likely to be generated from abundant and/or degradation-susceptible skin proteins we first defined an initial target cohort of 69 extracellular proteins (Fig. 2a and Table S1) drawn from the human skin proteome (Hibbert et al., 2018). In the case of a protease such as trypsin, which only cleaves at the C-terminal side of Lys and Arg (except when followed by Pro) (Manea et al., 2007), prediction of cleavage sites is straightforward. However, cleavage prediction for endogenous tissue proteases (such as MMPs and cathepsins) is determined by multiple factors other than primary amino acid sequence. Therefore, to identify the initial cohort of target proteins, we used an established machine-learning algorithm; PROSPER (Song et al., 2012) was used to predict the cleavage sites of skin-active enzymes (MMPs-2, -3, -7 and -9, cathepsins –G and –K, granzyme B and elastase-2 (Cavarra et al., 2002; Hiebert and Granville, 2012; Quan et al., 2013; Rijken et al., 2005; Xu et al., 2014)) for all human skin ECM proteins, and hence to select the 20 extracellular proteins with the highest proportion of predicted cleavage sites. As photo-oxidation can also degrade ECM components in skin (Berlett and Stadtman, 1997; Fisher et al., 1996; Wells et al., 2015), we next identified the 20 extracellular human skin proteins with the highest proportion of UVR/ROS susceptible amino acid residues (Hibbert et al., 2015). Finally, the cohort was supplemented with a further 33 ECM proteins (including collagens, elastic fibre associated proteins and proteoglycans) with known roles in dermal function. In order to reduce the computational requirements (of predicting multiple cleavage sites and potential matrikines), each protein in the initial cohort of 69 was further reviewed to select proteins with high relative abundance (immune staining on Human Protein Atlas (Hibbert et al., 2018) (Ponten et al., 2008)) and susceptibility to reported age-related remodelling. The final cohort of 27 target proteins included key components of the elastic fibre, system (i.e. elastin, fibrillin-1 and fibulin-1), fibrillar collagens (I and III) and the dermal-epidermal junction basement membrane (collagen IV and laminin-332) which all undergo remodelling in photoaged skin (Fig. 2a).

**Figure 2.**
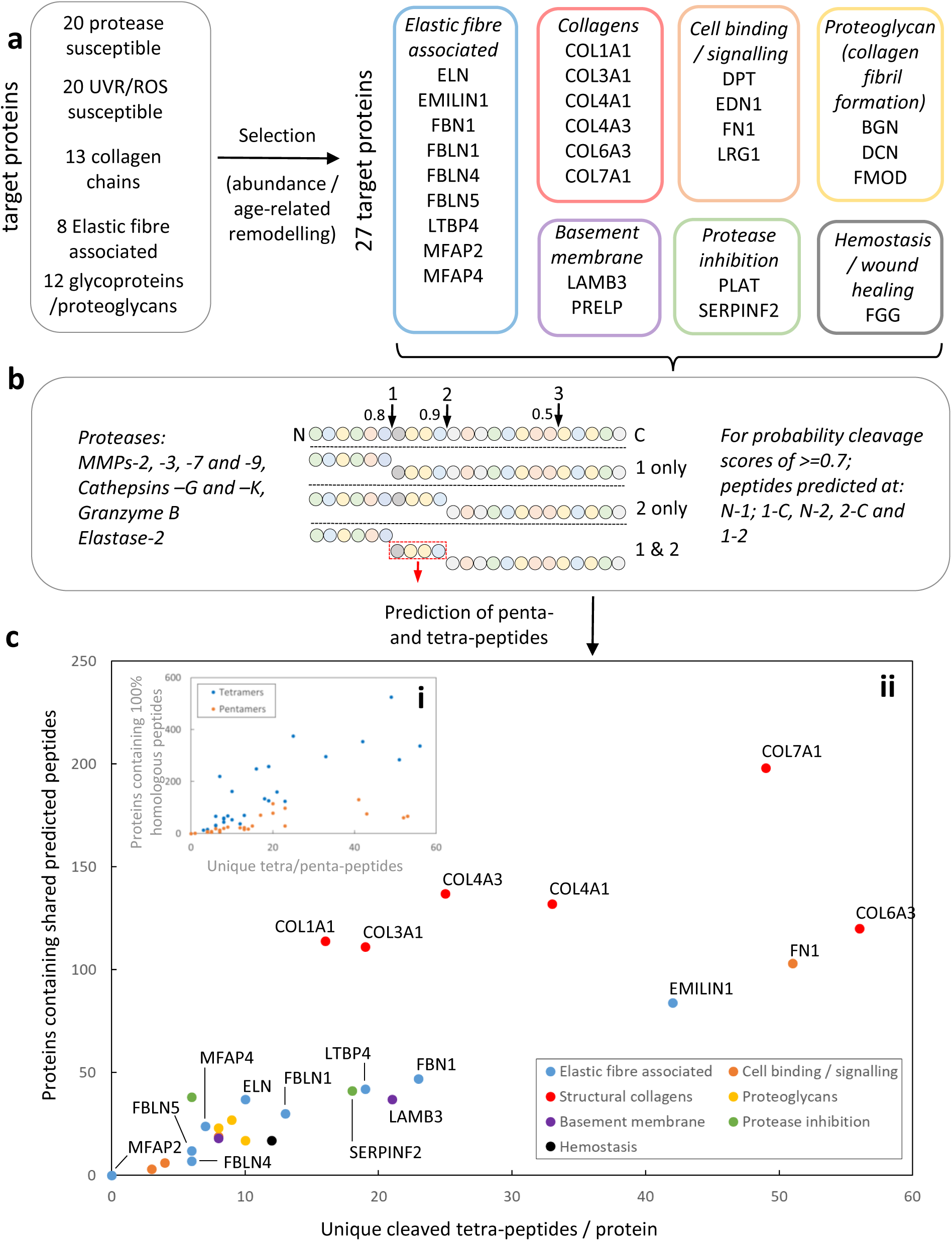
Bioinformatic prediction and selection of candidate peptides. (a) Defining protein targets for *in silico* protease cleavage. An initial cohort of 69 proteins (unique protease and UVR/ROS-susceptible and key structural ECM components) was filtered to identify 27 abundant and/or skin ageing-susceptible target proteins classified into seven categories. (b) *In silico* prediction of protease cleavage sites and liberated small peptides. Each of the 27 proteins was screened using a bespoke Python algorithm interacting with the PROSPER protease cleavage server to identify cleavage sites with a probability cleavage scores of >=0.7. In this hypothetical example a single tetra peptide (red dotted box) would be liberated. (ci) Comparable numbers of penta- (429) and tetra- (482) peptides were predicted to be cleaved from the target cohort but penta-peptide homologues (orange) were less widely distributed in skin proteins. (cii) For each protein, the number of unique cleaved tetra-peptides is plotted against the number of proteins which share predicted cleaved peptides with that protein. For example, COL7A1 harbours 49 predicted cleaved matrikines, which, collectively, are predicted to be cleaved from 198 skin proteins. In contrast, ELN contains 10 unique cleaved peptides, which are shared with 37 other proteins. In general, structural collagens, followed by some elastic fibre associated proteins and fibronectin, were predicted to be the most likely source of tetra-peptides. Collectively this screening process identified 453 unique putative matrikines.

### Prediction and *in vitro* selection of candidate matrikines from target proteins

Most ECM proteins are susceptible to digestion by multiple proteases and, as a consequence, have many experimentally-validated cleavage sites (Stewart-McGuinness et al., 2022). Therefore, we developed a bespoke algorithm which utilised PROSPER (Song et al., 2012) cleavage probability scores to predict fragmentation and hence the sequence of liberated peptides (Fig. 2b). Having generated a cohort of putative peptide fragments we next selected tetra-peptides for further review, as such small molecules (∼500Da or less) are more likely to penetrate the skin barrier (Bos and Meinardi, 2000) and may potentially be generated from 10’s-100’s of proteins. In contrast, penta-peptide homologues are less widely distributed in the skin proteome (Fig. 2ci), whilst di- and tri-peptides homologues are found in 1000’s of human proteins and hence may lack specific matrikine activities (Jariwala et al., 2022). Collagens IV, VI and VII are potentially rich sources of liberated matrikines, with between 25-56 unique cleaved tetra-peptide sequences predicted for the alpha chains of COL4A1, COL4A3, COL6A3, COL7A1 (Fig. 2cii). Not only do these collagens (along with COL1A1 and COL3A1) contain multiple predicted cleavage products but, for each alpha chain, the putative matrikines are also predicted (collectively) to be cleaved from over 100 skin proteins, thereby increasing the probability of them being liberated *in vivo*. In addition to the collagens, elastic fibre associated proteins (e.g. EMILIN1, FBN1, LTPB4), the adhesive glycoprotein FN1, and the basement membrane component LAMB3 may also be rich sources of matrikines *in vivo* (Fig. 2cii). This screening process identified 453 putative matrikine tetra-peptides.

Ubiquitous peptides may be produced with a higher frequency in damaged tissues, resulting in the promotion of ECM synthesis and tissue repair via multiple pathways. When selecting candidates for synthesis and experimental characterisation, we chose some peptides which were predicted to be cleaved from multiple skin ECM proteins by several proteases (in order to increase the chances of observing biological activity and/or broad-spectrum activity). For example, peptide P1 was predicted to be cleaved from multiple collagen alpha chains as well as ECM glycoproteins and ECM regulators. However, we also included some peptides which were predicted to be cleaved from a smaller and alternative protein cohort (for example P8 which is found predominantly in proteoglycans; Fig. 3a and Table S2). Additional review identified eight (8) peptides (P1: GPKG; P2: GPSG; P3: LSPG; P4: EKGD; P5: QTAV; P6: LSPD; P7: LSVD and P8: ELED) with predicted high solubility, the potential to form hydrogen bonds (which can mediate receptor/ligand interactions), and minimal issues with regards to downstream bulk-scale manufacture (Table S2). These were successfully synthesised and chemically modified with a palmitoyl chain (pal-) to increase penetration into the skin.

**Figure 3.**
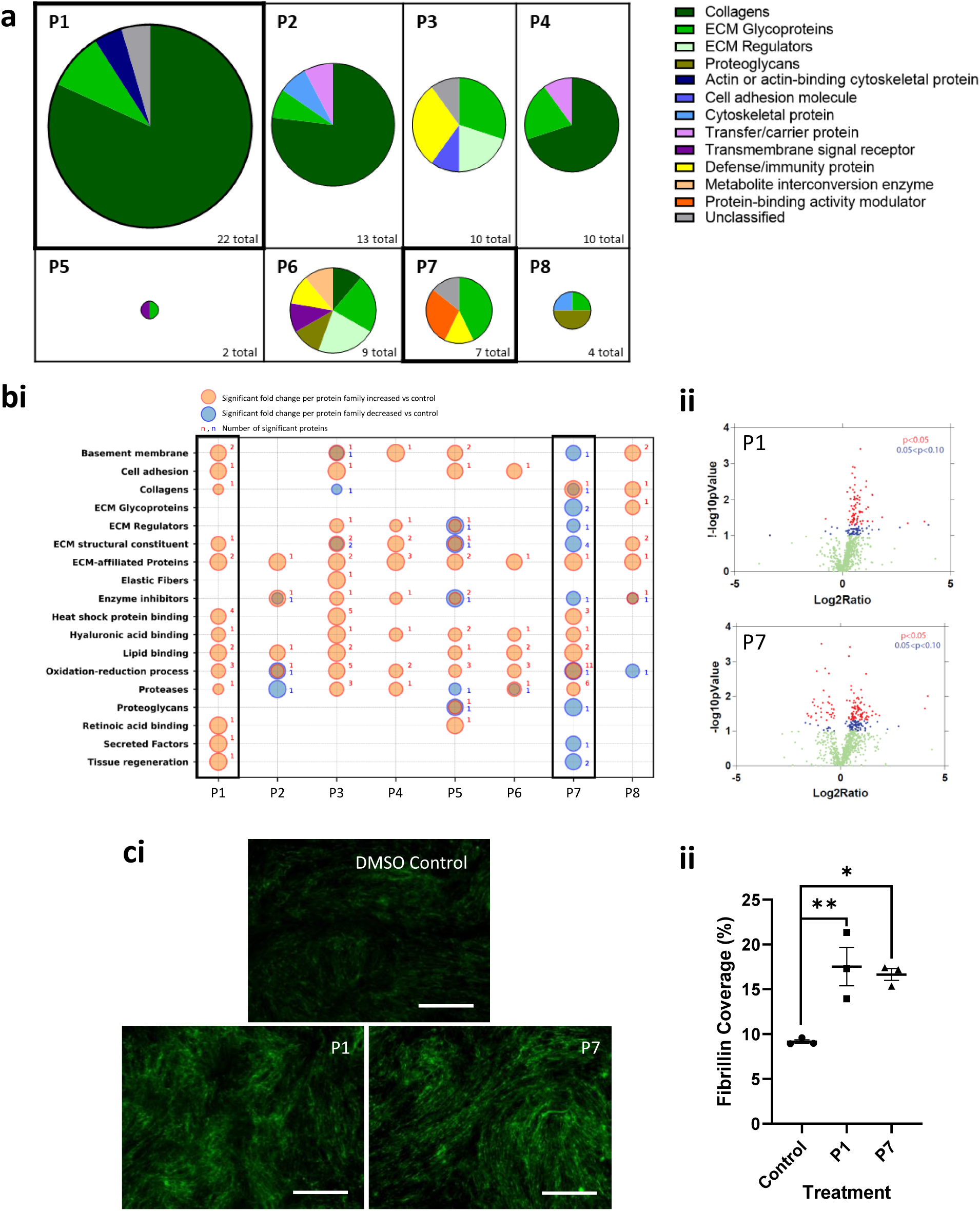
Characterisation of eight potential matrikine peptides. (a) Potential skin protein sources of peptide 1-8. Peptides P1, P2, P3, P4, P5, P6, P7 and P8 were originally identified as putative cleavage products of collagen I, emilin-1, collagen XVII, fibronectin, collagen VI, fibulin-1 and biglycan respectively. However, these peptides also have 100% homology to sequences found in many other skin proteins. Pie charts are sized relative to the number of proteins from which each peptide is predicted to be cleaved (i.e. 22 for P1 and 2 for P5). (b) Influence of peptides on the cultured HDF proteome. Cells were incubated for seven days either without any active added (untreated control), or with each peptide (n= 3 experimental replicates) for analysis of the combined secretome and cell matrix by LC-MS/MS. (bi) All peptides exhibited some ability to influence the cell with P1, P3, P5 and P7 showing the broadest range of activity. P1 significantly enhanced protein synthesis in wide range of protein families (bi and bii volcano plot red points). Whilst P7 induced both significant up- and down-regulation of proteins (bi and bii volcano plot). The identity of proteins up- and down-regulated by P1 and P7 is reported in the supplementary data. (c) Induction of fibrillin-1 synthesis in cultured HDFs (at optimum concentration for each peptide) following 5 days treatment (n = 3 experimental replicates). Both P1 and P7 peptides enhanced deposition of a fibrillin-rich microfibrils network (green stained filaments in ci) compared with the DMSO-treated control Scale bar = 100μm. Fibrillin rich-microfibril deposition was significantly higher for P1 (adjusted p = 0.0069) and P7 (adjusted p = 0.0117) by one-way ANOVA using Dunnett correction for multiple comparisons (cii).

### Selection of two peptides with promising *in vitro* activities

Initial toxicity testing (Hoechst 33258) was conducted to determine viable peptide concentrations for *in vitro* testing (Table S3). The ability of the 8 peptides to promote cellular synthesis of selected ECM components essential to dermal integrity (procollagen I, fibronectin, decorin, collagen IV, hyaluronic acid and fibrillin-1) was then assessed *in vitro* on human dermal fibroblast cells (HDFs) via enzyme-linked immunosorbent assays (ELISA) or immunofluorescence (IF) techniques. With the exception of hyaluronic acid, all peptides enhanced the synthesis of at least some of the ECM markers tested (Table S4). These targeted immune-assays were followed by liquid chromatography tandem mass spectrometry (LC-MS/MS) proteomic analysis of HDFs exposed to each peptide. Although peptides P2, P4, P6 and P8 upregulated proteins in multiple functional classes (Fig. 3b), their activity was more limited than the other peptides and peptides P3 and P5 were present in few skin proteins. Consequently, two peptides with contrasting activities (P1 and P7) were selected for characterisation of fibrillin-rich microfibril (an early biomarker of both skin ageing and repair (Watson et al., 2008)) deposition. Both peptides induced significant elaboration of a fibrillin-rich microfibril network compared with the DMSO control (Fig. 3ci). P1 was therefore selected for further characterisation due to its ubiquitous distribution within skin proteins and ability to enhance pro-collagen I synthesis (by ELISA) and a broad range of proteins, including fibrillin-rich microfibrils (IF) and basement membrane components (LC-MS/MS; Fig. 3cii). P7 was also chosen as it promoted both fibrillin-1 and decorin synthesis (by IF and ELISA respectively) and showed diverse activity in the LC-MS/MS screen.

### Peptides pal-GPKG (P1) and pal-LSVD (P7) act synergistically in vitro

Primary HDFs from five age- and sex-matched donors (derived from breast skin (n=1), labia (n=2) and buttock(n=2)) were treated with P1, P7, P1+P7 in combination and TGF-β1 (which acting as a positive control profoundly affected both the cellular transcriptome and proteome; Figure S1ai). Total RNA was extracted after 12 hrs (in order to characterise the initial response to peptides) and extracellular proteins after seven days of treatment (to characterise protein deposition). RNA-Seq transcriptome analysis identified over 6,000 genes with peptide-treatment associated fold changes of ≥1.2 (Yao et al., 2015) in each of the peptide treatments. Crucially, whilst principal component analysis (PCA) demonstrated strong clustering of samples by body site (Fig. 4ai), PCAs for each individual donor (Fig. 4aii and Figures S1ai-vi) showed that in all cases, peptide treatments modulated the transcriptome compared with untreated controls. Gene Ontology (GO) term enrichment analysis highlighted the diversity of biological processes enriched by the peptides individually (e.g. P1, cellular proliferation and lipid metabolism; P7, ECM remodelling). However, there was a profound synergistic effect, with the peptides in combination (hereby referred to as P1+P7) significantly enriching the transcription of 243 processes (compared with 84 for P1 and 52 for P7 alone). Of these; P1+P7 induced 32 processes involving ECM genes (as defined by the 27 ECM target genes listed in Table S1) compared with 6 for P1 and 11 for P7 (Fig. 4b and Figure S1). In particular, P1+P7 enriched processes concerned with ECM assembly and cell adhesion and with cellular proliferation. Proteomics analysis confirmed the transcriptomic clustering by HDF body site origin but identified more separation between untreated control cells and 7 days peptide treatments (Fig. 4c). Again, the peptide combination was most biologically active, upregulating the deposition of over 50 ECM proteins *in vitro* (as defined by Matrisome DB (Shao et al., 2020)) with a fold change >=1.2 (Fig. 4c and d). In particular, the combination enhanced synthesis of ECM regulators and proteoglycans, including serpin peptidase inhibitor, clade E, member 1, and versican core protein.

**Figure 4.**
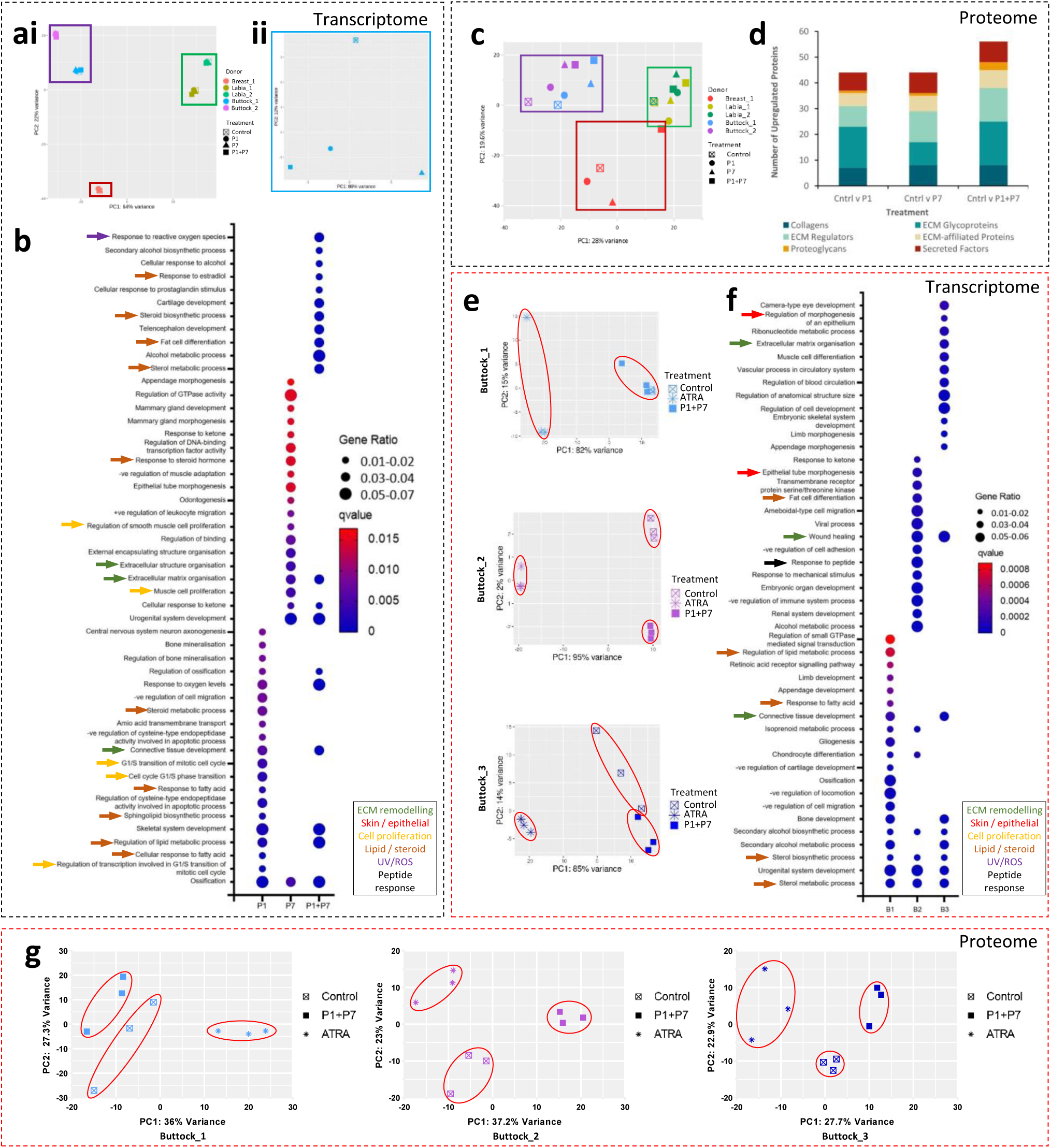
In vitro characterisation of the biological activities of peptides 1, 7 and the combination. (a-d) In vitro ‘omics characterisation of HDFs derived from multiple skin sites exposed to individual peptides and peptides in combination (n= 5 biological replicates per treatment). (ai) Principal component analysis (PCA) of RNA-Seq data for peptide exposed HDFs. Gene expression clustered by cell donor and by cell donor site (i). However, for each individual donor (aii: example PCA for buttock 1) there was clear separation between treatments. (b) GO-term enrichment analysis of RNA-Seq data for the top 20 (q-value) enriched biological processes. Both P1 and P7 treatments enriched multiple yet distinct processes but the peptides in combination acted synergistically to upregulate a wider range of processes with lower q values and higher gene ratios. (c) Proteomic analysis: PCA of LC-MS/MS data. After 7 days exposure to treatments, whilst protein synthesis remained clustered by skin site, the influence of the peptides was more readily discernible. (d) Upregulated ECM proteins with a fold change >=1.2. Of the 700 proteins upregulated by the peptide combination, over 50 were ECM components. (e-g) Characterisation of buttock-derived HDFs exposed to the peptide combination with ATRA as a positive control (n= 3 biological replicates with 3 experimental replicates per treatment). (e) PCAs of individual donor RNA-Seq data. Both ATRA and peptide treatments induced clear and distinct clustering compared to the control. (f) In the top 20 (q-value) enriched biological processes, sterol biosynthesis (yellow arrows) was enhanced in all three donors and ECM-related tissue development/repair processes in 2/3 donors. (g) PCAs of proteomic data. Peptide treatment induced clear clustering in all three donors, which was distinct from the control and ATRA treated cells.

Given the clear synergy between the peptides, the influence of body site on HDF phenotype, and the profound effects of TGF-β1 on cell phenotype, we next tested the biological activity of the peptide combination against solely buttock-derived primary HDFs (n=3) using a more clinically relevant positive control (all-*trans* retinoic acid; ATRA). The combination peptide treatment once again significantly enhanced transcription of genes relevant to ECM-rich tissues (ECM organisation, collagen biosynthesis, connective tissue development and wound healing) and lipid/steroid-related processes, but also to peptide response and epithelial processes (Fig. 4e and f and Fig. S2). After 7 days’ exposure, the peptide combination induced clear clustering by the proteins induced on the PCA plots, which was distinct from both the control and those treated with ATRA (Fig. 4g). Over 20 ECM proteins in each donor dataset were found to be upregulated by the peptide combination treatment, with ECM glycoproteins and regulators being in the majority. Notably, fibronectin, vitronectin, fibromodulin, as well as MMPs-1 and -14 were all found to be upregulated in multiple cell donors.

### The peptide combination acts *in vivo* to promote epithelial and dermal remodelling

Before the peptide combination could be progressed to *in vivo* efficacy testing on human skin, the peptides (500 ppm) were solubilised into a suitable and stable excipient for use in topical formulations and subjected to a range of *in vitro* and *ex vivo* toxicology assessments following Organisation for Economic Co-operation and Development (OECD) guidelines (Development). QSAR and Xenosite analysis, SkinEthic, Epiocular, Uvs spectra, DPRA and keratinosens tests were performed on two peptides (P1 and P7). The peptide formulation (containing 50:50 ratio of P1 and P7) was deemed to be neither a skin sensitizer nor an eye irritant and could, therefore, be safely progressed to *in vivo* efficacy assessment.

Chronic UVR exposure results in epidermal thinning and significant remodelling of the underlying dermal ECM, including loss of fibrillin-rich microfibrils (oxytalan fibres) from the superficial papillary dermis and accumulation of dystrophic elastin (solar elastosis) in both the papillary and reticular dermis (Naylor et al., 2011). To determine the efficacy of peptides P1 and P7 in mitigating epidermal and dermal remodelling, the peptide formulation was applied to the photoaged extensor forearm skin of otherwise healthy volunteers, using a validated, occluded patch test assay (Watson et al., 2008) alongside an occluded but untreated area (occluded control), a vehicle control and ATRA (as positive control). In contrast to *in vitro* characterisation, which employed a fibroblast monoculture, the *in vivo* patch test exposed multiple cell types embedded in a complex tissue environment to the peptide combination. In the bulk RNA-Seq analysis, for all donors, the peptide formulation modulated the transcriptome compared with both the occluded and vehicle controls (Fig. 5a). Demonstrating that the peptides exert a significant influence on cell physiology additional to the vehicle. Specifically, GO-term analysis of RNA-Seq data for the top 20 (by q-value) biological processes identified multiple differentially enriched processes against the vehicle control: at 10ppm these included peptide and peptide hormone responses and retinol metabolism. At 30ppm, against the vehicle control, the peptide formulation drove expression of key skin homeostasis and repair processes, including keratinization, cornification, and skin barrier (Fig. 5b). Compared with the occluded control transcriptome, the peptide formulation enhanced transcription of multiple ROS- and ECM-related processes (Fig, 5c). In addition to modulating expression of collagens, laminins and proteoglycans, the peptide formulations enriched transcription of elastic fibre components, including elastin, fibulins and microfibril associated protein 4 (MFAP4). Immunohistochemical analysis demonstrated that compared with the occluded control, the peptide formulation significantly enhanced deposition of fibrillin-rich microfibrils in the papillary dermis (Fig. 6a and b). However, the vehicle control did not significantly affect fibrillin-rich microfibril deposition compared with the occluded control, indicating the role played by the peptides in stimulating elastic fibre remodelling.

**Figure 5.**
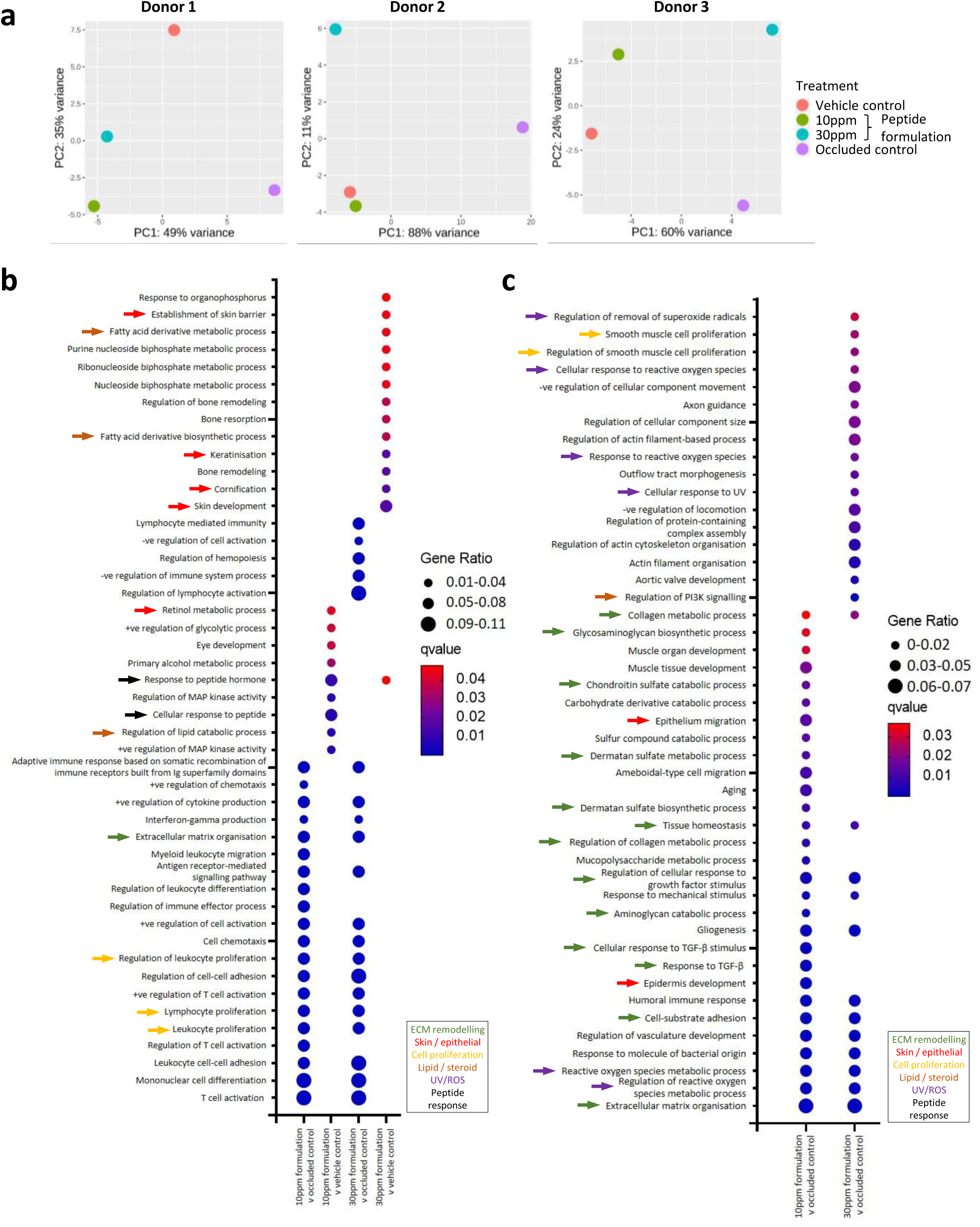
In vivo transcriptomic characterisation of peptide treated skin. (a) PCA of RNA-Seq data for three individual donors: occluded control, vehicle (control), ATRA (positive control) and P1+P7 at concentrations of 10ppm and 30ppm. For all three donors ATRA treated transcriptomes segregated according to PC1. Peptide treated transcriptomes segregated from the occluded control and (with the exception of donor 2) the vehicle, by PC2. (b) GO-term analysis of RNA-Seq data for the top 20 (q-value) enriched biological processes for peptides at both concentrations vs occluded and vehicle controls. Topically applied peptides preferentially enriched immune cell proliferation (occluded control: yellow arrows) and skin remodelling (vehicle: red arrows) processes. (c) GO-term enrichment analysis for ECM genes of interest against the occluded control (27 genes defined in Extended Data Table 1: collagens, elastic fibre associated and adhesive glycoproteins and proteoglycans). At both concentrations, the peptide combination enriched multiple ECM remodelling processes (green arrows). Whilst ROS- and UV-related processes were predominantly enriched by 30 ppm P1+P7.

**Figure 6.**
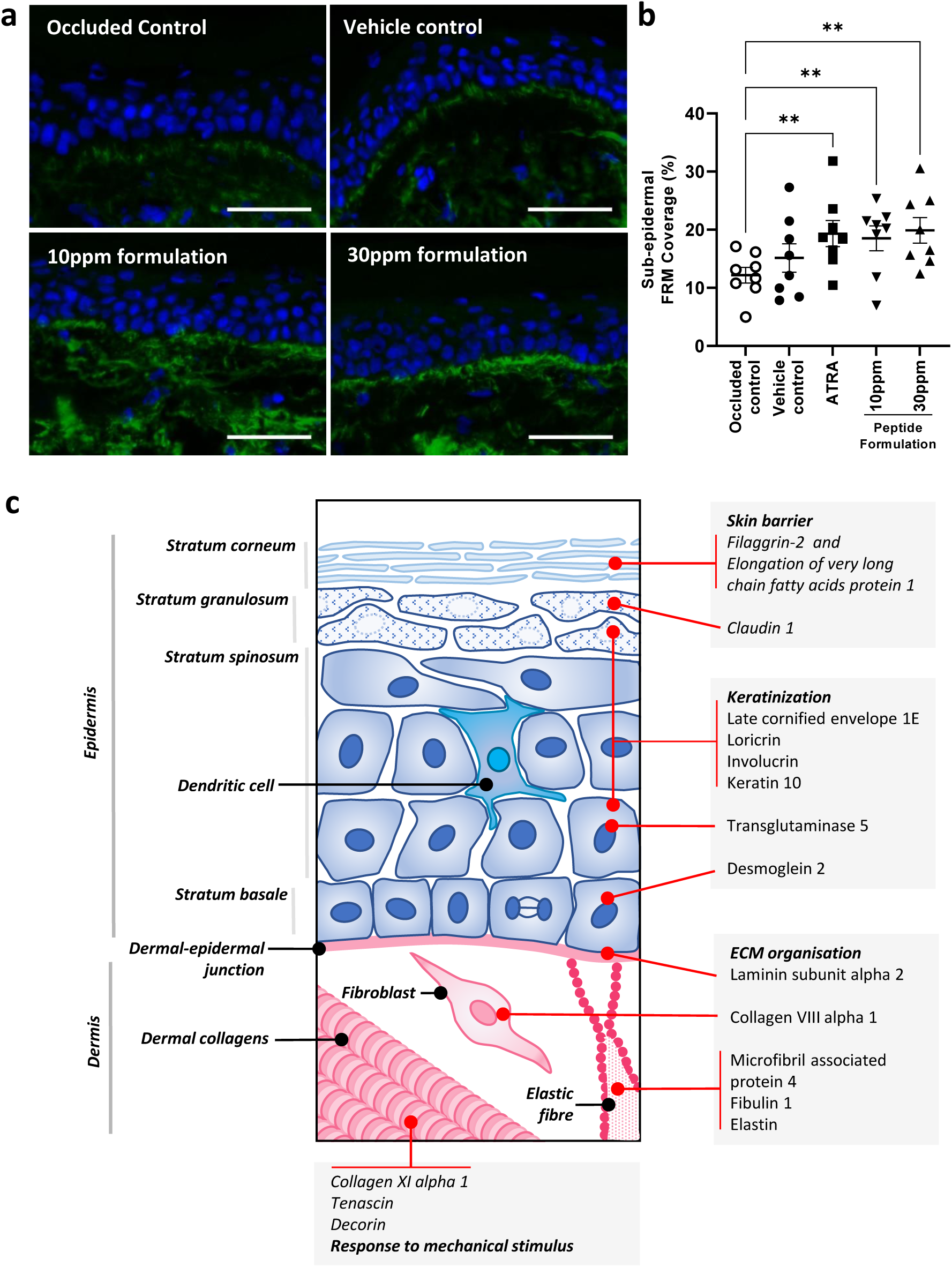
Peptide-treated skin: immunohistological characterisation and impact on biological processes. (**a**) Immunofluorescence staining of fibrillin-rich microfibrils, following 12-day patch test with occluded control, vehicle and 10ppm and 30ppm peptide blends. A 4–day occluded ATRA patch test was included as a positive control (n = 8 biological replicates per treatment). (b) Analysis of immunofluorescence of fibrillin-rich microfibrils. Sub-epidermal microfibril coverage was measured as a percentage of area covered with microfibrils within an area 30μm immediately below the epidermis. Repeated Measures one-way ANOVA was used for statistical analysis with where significance *p < 0.05 **p< 0.005. Error bars represent mean ± SEM (n=8). (c) Exposure to the peptide formulation enhances biological processes key to skin health including the epidermal maturation of keratinocytes and development of a skin barrier and the expression of dermal ECM components involved in elastic fibre deposition and modulation of tissue biomechanical properties.

## Discussion

In this study we test the hypothesis that small bioactive peptides (matrikines) can be predicted by the *in silico* digestion of dermal ECM proteins by proteases. Our discovery pipeline: i) provides *in vitro* evidence for diverse, sequence-dependent, biological activities induced in cultured HDFs exposed to exogenous tetra-peptides and; ii) shows that the two peptides tested *in vivo*, when applied in combination, can modulate key measures of skin photo-damage. Although the detection of peptides less than 500 Da in a complex proteome is challenging with conventional mass spectrometry, more targeted approaches such as selective or multiple reaction monitoring, could be used to identify previously predicted small peptide sequences in complex protein mixtures extracted from growing or healing tissues. Although there is evidence that some ECM fragments (such as collagen XVIII-derived endostatin (Isobe et al., 2010)) can reduce fibrosis, the role played by endogenous matrikines in mediating self-repair in ageing and/or diseased tissues is an important area for study. The complex nature of ageing and many chronic diseases, where a diverse array of proteases may act on hundreds of proteins to produce thousands of peptides at low concentrations, could explain the lack of evidence for endogenous matrikines in preventing tissue degeneration. Even with a small cohort of ECM derived peptides, our data demonstrates that each sequence (peptides P1-P8) can mediate the expression of numerous and disparate proteins and pathways. By applying relatively high concentrations of a peptide, the cells receive a strong and consistent signal, in contrast to the potential low-level noise of many endogenously derived peptides.

Predicting and selecting peptides with specific activities will be challenging. Whilst our data indicates that some peptides (i.e. P3) may act as relatively specific alarm signals prompting cells to primarily synthesise enzyme inhibitors (Fig. 3c). Other peptides (particularly P1, P3, P5 and P7) appear to be non-specific alarm signals, inducing fibroblasts to synthesise a large and varied proteome (Fig.3c). The limited nature of outcome measures assessed for commercially-available skin-active peptides makes it difficult to make comparisons with the transcriptomic and proteomic measures used in this study, but our data confirms previous observations (Jariwala et al., 2022) that ECM-derived peptides promote ECM synthesis. It is possible that longer (and hence protein source-specific) peptide sequences may have more targeted effects but, for use in topical skin treatments, penetration through the *stratum corneum* may be challenging (Jariwala et al., 2022) and smaller peptides (Cans (Chamani and Zamani, 2022)) can exhibit similar activities to the parent molecule (canstatin (Kamphaus et al., 2000)). Relative peptide activity will also be subject to inter-individual variation (Figs. 4-6). In the case of skin, it is well established that even the gold-standard topical treatment ATRA varies in its effectiveness between individuals (Kligman et al., 1986; Watson et al., 2008; Young et al., 2006). The biological effects of exogenous peptides will also be target-cell dependent and therefore single cell transcriptomic and proteomic profiling followed by expression quantitative trait loci (eQTL) analysis on well-characterised cell lines such as HipSci fibroblast lines may provide insight towards personalised treatment approaches(Kumasaka et al., 2021; McCarthy et al., 2020). In common with most previous studies, we have used cultured dermal fibroblasts as the target cell to detect and respond to peptides. It is clear that fibroblasts are responsive to the exogenous peptides, but *in vivo* epidermal cells, and in particular keratinocytes, will be exposed to the highest doses. Our *in vivo* patch test protocol and subsequent transcriptomic characterisation highlights the enrichment of key epithelial processes including skin barrier formation, keratinization and cornification (Hussain and Goldberg, 2007; Puig et al., 2008) indicating that the effects of this peptide combination are not confined to the dermis.

By applying an *in silico* to *in vivo* discovery pipeline we have identified and characterised the ability of novel peptides (GPKG and LSVD) to enhance the transcription of ECM organisation and cell proliferation genes and to promote epithelial and dermal remodelling (Fig. 6c). The use of such biomimicry approaches to predict the identity of naturally occurring ECM breakdown products which promote cell signalling could facilitate the development of safe and well-tolerated technologies and therapies. The development of improved techniques for predicting and detecting the cleavage of small peptides and for the localisation of peptide action within organs and target cells will be critical to enabling better understanding of the mechanisms of matrikine-induced tissue repair with both patient and consumer benefits.

## Materials and Methods

### Discovery pipeline overview

Beginning with bioinformatic screening of the human skin proteome, the discovery pipeline integrated existing resources (PROSPER protease cleavage prediction server (Song et al., 2012)) with bespoke algorithms to predict cleavage sites and hence potential peptide fragments from 27 target proteins) with matrikine activity. Eight candidate matrikines, selected on the basis of predicted cleavage from multiple source proteins, reported literature involvement in skin ageing and assessments of suitability for manufacturing, were synthesised, modified with a palmitoyl chain and characterised for biological activity in cultured HDFs. Following initial immunological screening with key ECM components as outcome markers, the biological and toxicological effects of two candidate peptides (individually and in combination) were assessed. The peptide combination was subsequently applied in a formulation to the skin of eight human volunteers and the impact on skin histology and transcriptome characterised.

Using an adapted systematic review approach, we previously defined the human skin proteome, which includes 205 constituent dermal ECM proteins (Hibbert et al., 2018) (Fig. 1a). In order to identity dermal ECM proteins for matrikine prediction, we first defined a cohort comprising protease- and UVR/ROS-susceptible proteins supplemented with further ECM structural proteins. The 20 most protease-susceptible extracellular proteins within the human skin proteome were predicted by *in silico* digestion (PROSPER) by MMPs-2, -3, -7 and -9, cathepsins –G and –K, granzyme B and elastase-2 (Cavarra et al., 2002; Hiebert and Granville, 2012; Quan et al., 2013; Rijken et al., 2005; Xu et al., 2014). PROSPER uses publicly available data from the MEROPS database including manually curated and experimentally validated protease cleavage consensus sequences (Rawlings et al., 2018). Using a support vector machine learning (SVM) algorithm (Joachims, 1999) PROSPER estimates the likelihood of proteolytic cleavages occurring in particular positions on protein primary sequences. To make predictions more robust, the PROSPER classifier integrates structural properties such as disordered regions (regions that are capable of undergoing structural changes while performing their functions (Uversky, 2013)) as predicted by DISOPRED3 (Jones and Cozzetto, 2015), assessments of protein secondary structure (psipred (Buchan et al., 2013)) and solvent accessibility (ACCpro 4.0 (Cheng et al., 2005)). The 20 most protease-susceptible proteins were defined as those with the greatest number of cleavage sites (with a cleavage probability score (Song et al., 2012) of >=0.9) per protein length. Next, having previously shown that amino acid composition is an important factor in determining the susceptibly of proteins to photodynamic degradation (Hibbert et al., 2015) we calculated the relative proportions of oxidation and UVR-sensitive amino acid residues (Trp, Tyr, His, Met, Cys and Cys=Cys) in each extracellular human skin protein to define a further 20 proteins most at risk of photodynamic degradation. A final tranche of structural collagens (each constituent alpha chain analysed separately), elastic fibre associated proteins and adhesive glycoproteins and proteoglycans was added to define an initial cohort of 69 proteins (taking into account proteins identified in more than one category) (Table S1). Following further review of these proteins we identified a sub-cohort of 27 target proteins (Fig. 2a) with relatively high skin abundance (as assessed by immune staining in the Human Protein Atlas (Ponten et al., 2008), and susceptibility to age-related remodelling (as reviewed in the Manchester Skin Proteome (Hibbert et al., 2018)).

### *In silico* peptide prediction

In order to predict the identity of cleavage sites and hence cleavage products for each of the 27 proteins, protein FASTA sequences along with domain information were retrieved from Uniprot and the sequences were digested *in silico* by the same panel of enzymes as used in target protein selection (MMPs-2, -3, -7 and -9, cathepsins –G and –K, granzyme B and elastase-2). We developed a bespoke algorithm (https://github.com/maxozo/Matrikine_Discovery) which to predict the sequence of liberated peptides. This algorithm utilised the PROSPER predictions of protease cleavage sites to predict cleavage sites and hence peptide fragments (where the average N- and C-terminal cleavage probability score was >= 0.7) (Fig. 2b). This algorithm was also applied to all remaining proteins in the human skin proteome. Following a homology comparison of predicted tetra-peptide sequences with predicted protein fragments for all human skin proteins, 453 peptides from the 27 target proteins were selected for further review. A solubility score (average hydrophilicity of the constituent amino acid residues) was then calculated for each peptide (Hopp and Woods, 1981) as well as a hydrogen bond score (score of 1 for amino acid residues able to form hydrogen bonds [Met, Lys, Thr, Asp, Glu, Ser, Cys, Tyr, His, Asn, Gln, Arg, Trp] and score of 0 for the remaining residues) (Barrett, 2012). Additionally, a “Potential Problem Score” was assigned to each peptide based on the presence of amino acids susceptible to oxidation, vicinity of highly hydrophobic amino acids and or to avoid side reactions during synthesis or formulation.

A cohort of eight peptides was selected for synthesis and for screening. These peptides were predicted to be highly soluble, ability to form hydrogen bonds and to have few potential manufacturing problems with non CMR solvents. The cohort contained peptides which were predicted to be cleaved from a wide range of ECM components (Fig. 3a).

### Synthesis and toxicity testing of eight candidate peptides

All eight peptides were successfully synthesised and chemically modified with a palmitoyl chain to increase penetration into the skin (Choi et al., 2014). Peptides were synthesized via a solid phase synthesis with sequential coupling of Fmoc-amino acids from C-terminal to N-terminal, then, coupling of palmitic acid. Synthesis was performed without any CMR solvents. Pure peptides were obtained with a purity of 99%, assessed by high-performance liquid chromatography–mass spectrometry (HPLC-MS; mass detector 6120 quadrupole, HPLC-1200; both from Agilent, USA).

*In vitro* toxicity was assessed for each peptide using cultured primary HDFs from juvenile foreskin (Celln Tech™) and the Hoechst 33258 toxicity assay (Labarca and Paigen, 1980) (Table S3). HDFs were cultured for 72 hours in the presence of each peptide (at 3-12.5 ppm)in Dulbecco’s modified Eagle’s medium (DMEM, Gibco #21969-035) supplemented with 10% fetal bovine serum (FBS, Gibco #10270-098), 100 U/mL penicillin, 100 mg/mL streptomycin (Gibco #15070-063), 1μg/mL fungizone (Gibco #15290-026) and 1mM L-glutamine (Gibco #25030-024). HDFs were incubated at 37°C in a humidified 5% CO2 atmosphere. Cells were rinsed with phosphate-buffered saline (PBS; Gibco #20012027), then, an ultrasound-induced cell lysis was performed. Cell DNA contents were stained using 8μg/mL Hoechst 33258 solution in PBS (Sigma #B2883) for 30 min. Cell number evaluation was performed with BMG Omega fluostar (ex/em. 360/460nm) following Labarca and Paigen method allowing the cell viability when exposed to each peptide to be calculated.

### Initial activity screens of eight candidate peptides by ELISA, fibrillin-I immunofluorescence and LC-MS/MS proteomics

Peptide activity was initially tested by characterising the ability of each peptide to induce synthesis of a panel of dermal ECM markers: pro-collagen I, fibronectin, decorin and collagen IV measured by ELISA (n=2 with five experimental replicates for all proteins apart from collagen IV with four experimental replicates). Primary HDFs from juvenile foreskin (CellInTech^TM)^) were cultured in routine 175cm^2^ flask (Falcon™) with Dulbecco’s modified Eagle’s medium (DMEM, Gibco #21969-035), 10% FBS (Gibco #10270-098), 100 U/mL penicillin, 100 μg/mL streptomycin (Gibco #15070-063), 1μg/mL fungizone (Gibco #15290-026) and 1mM L-glutamine (Gibco #25030-024), under 5 % CO_2_ and 90% humidity atmosphere at 37°C. Medium was renewed every 48h to 72h to allow outgrowth. HDFs were used between passage 5 and 11. Thereafter, HDF cultures were seeded at a density of 2 × 10^4^ cells / cm^2^ in 24 well cell culture plates (Falcon #353226) and were treated with various concentrations of peptides (as determined by prior Hoechst 33258 staining in serum–free media) or solvent (DMSO, 0.1% v/v, Merck) or positive control TGF-β1 10ng/ml (Sigma, T7039) for 3 days in fresh medium (without FBS due to assay interference). Then, cell culture media were collected for proteins and glycosaminoglycan release measurements using ELISA methods; procollagen type I (Takara #MK101), fibronectin (Takara #MK115), collagen IV (Magnetic Luminex Assay Collagen-IV, R&D System #LXSAHM-1), hyaluronic acid (Corgenix 029-001) and decorin (R&D Systems DY143) assays respectively. ELISA assays were performed as per the manufacturer’s instructions. All treatments were run in pentaplicate or tetraplicate (collagen IV) wells per experiment. In parallel, cell viability was estimated using Hoechst 33258 staining (Sigma #B2883) as per Labarca and Paigen’s (1980) protocol allowing the quantity of dermal proteins to be weighted to the number of viable cells. One-way analysis of variance (ANOVA) was used to determine whether there was any significant difference between the means of two or more independent groups in the ELISA assays, with p-values of *<0*.*05(*\*) or *<0*.*01*(**) considered statistically significant. Difference between two means with similar variances was performed with Student’s t-test.

The effects of each candidate peptide on fibrillin-1 synthesis were assessed by immunofluorescence. Primary HDFs from abdominal skin of an adult, Caucasian 29-year-old female (Promocell) were cultured in HDF growth media supplemented with FBS (2% v/v) basic fibroblast growth factor (recombinant human; 1ng/ml) and insulin (5 μg/ml) (all from PromoCell). All cells were grown in a humidified environment at 37°C with 5% CO_2_. For treatment and fibrillin IF, cells at passage 3-4 were seeded in to black walled clear bottomed 96 well plates (Pierce, ThermoFisher Scientific) at a density of 7,500 cells per well. After 24 hours, media was removed and cells were cultured for 5 days in the presence of DMSO (0.1% v/v)-supplemented media alone or media supplemented with the novel tetra-peptides solubilised in DMSO (0.1% v/v) at varying concentrations (as determined by prior Hoechst 33258 staining in serum-containing media), changing media every 48 hours. All treatments were run in triplicate wells per experiment. After 5 days, the media was discarded, and cells and surrounding ECM fixed with ice-cold methanol at -20°C for 5 minutes. Following PBS washing and blocking with 5% bovine serum albumin in PBS for 30 minutes, cells were then probed using a primary antibody to extracted fibrillin-rich microfibrils (11C1.3; mouse monoclonal antibody; ThermoFisher Scientific; 1 in 100 dilution overnight) designed to detect intact fibrillin-rich microfibrils, followed by a secondary goat anti-mouse IgG H&L Alexa Fluor 488 (Abcam; 1:1000 dilution). Fluorescence was then visualised using an Eclipse 100 microscope (Nikon, Japan).. Images were taken at a set exposure time Image-J software was used to quantify the fibrillin-1 coverage (%) in each image using a set threshold per experiment. To test for statistical significance, a repeated measures one-way ANOVA using Dunnett correction for multiple comparisons was performed, where adjusted p-values of *<0*.*05(*\*) or *<0*.*01*(**) considered statistically significant.

The impact of each candidate peptide on the HDF proteome as assessed by LC-MS/MS. Primary HDFs from abdominal skin of adult, Caucasian females aged between 23-33 years were cultured in Dulbecco’s Modified Eagle Medium (DMEM) with 4.5 g/l D-Glucose and L-Glutamine supplemented with 10% FBS, 100 μg/ml penicillin, 100 μg/ml streptomycin and 2 mM Gibco^®^ GlutaMAX™ (an alternative L-glutamine supplement). All cells were grown in a humidified environment at 37°C with 5% CO_2_. Cell media (secretome) was collected and the matrix decellularised (using EDTA). Secretome samples were added to the decellularised ECM left behind in the plate and proteins were denatured (in urea), reduced (in dithiothreitol), alkylated (in iodoacetamide) and mechanically disassociated (using ultrasonication) prior to overnight digestion with the SMART Digest kit as per optimised protocol. Samples were run on a an UltiMate® 3000 Rapid Separation Liquid Chromatographer coupled to a Q Exactive™ Hybrid Quadrupole-Orbitrap™ Mass Spectrometer in DDA mode (200 ng injected at 90-minute runs per sample).

### In vitro characterisation of cell transcriptomes in response to peptides 1, 7 and 1+7 in combination

The effects of the two candidate peptides was assessed by RNA-Seq analysis. Primary HDFs from adult Caucasian females aged between 23-33 (taken from a range of skin sites: buttock (n=3), labia (n=2) and breast (n=1)) were seeded within appropriate cell densities into tissue culture plates in assay medium – Fibroblast Basal medium-2 (phenol red-free; PromoCell, C-23225) containing 2% FBS, 0.001% basic human fibroblast growth factor (hbFGF) and 0.005% insulin (Fibroblast growth medium 2 supplement pack; PromoCell, C39320)). After two days of culture in a humidified environment at 37°C with 5% CO_2_, once cells had reached confluency, cells were treated. Treatments included 8 parts per million (ppm) P1, 8ppm P7, a combination of P1 and P7 at 8 ppm each, 0.2 nM TGF-β1 (Gibco, PHG9214) and 1 μM ATRA (Sigma-Aldrich, R2625).

For analysis of cellular transcriptomes, RNA was extracted 12 hours following the treatment using the RNeasy Mini Kits (74104, Qiagen) according to the manufacturer’s instructions. Briefly, lysis buffer was added to the samples, followed by precipitation of nucleic acids and desalting by addition of 70% ethanol. The resulting mixture was added to the RNeasy Mini Spin Columns and centrifuged to bind total RNA to the membrane. Samples were then washed, first with buffer RW1 to remove biomolecules such as carbohydrates, proteins and fatty acids which may have bound non-specifically to the membrane. The membrane was then washed twice with buffer RPE to remove salts from previous steps of the method. Finally, using RNAse-free water, total RNA was eluted from the spin columns. All extraction steps were performed on ice, with pre-chilled solutions. Following RNA extraction, RNA sample concentration was measured using a NanoDrop™ Spectrophotometer (ThermoFisher Scientific). All samples were stored at -80°C, and repetitive freeze-thaw cycles were avoided by aliquoting RNA samples. RNA samples were analysed as described below.

Total RNA was submitted to the Genomic Technologies Core Facility, Faculty of Biology, Medicine and Health, University of Manchester. Quality and integrity of the RNA samples were assessed using a 4200 TapeStation (Agilent Technologies) and then libraries generated using the Illumina® Stranded mRNA Prep. Ligation kit (Illumina, Inc.) according to the manufacturer’s protocol. Briefly, total RNA (typically 0.025-1ug) was used as input material from which polyadenylated mRNA was purified using poly-T oligo-attached, magnetic beads. Next, the mRNA was fragmented under elevated temperature and then reverse transcribed into first strand cDNA using random hexamer primers and in the presence of Actinomycin D (thus improving strand specificity whilst mitigating spurious DNA-dependent synthesis). Following removal of the template RNA, second strand cDNA was then synthesized to yield blunt-ended, double-stranded cDNA fragments. Strand specificity was maintained by the incorporation of deoxyuridine triphosphate (dUTP) in place of dTTP to quench the second strand during subsequent amplification. Following a single adenine (A) base addition, adapters with a corresponding, complementary thymine (T) overhang were ligated to the cDNA fragments. Pre-index anchors were then ligated to the ends of the double-stranded cDNA fragments to prepare them for dual indexing. A subsequent PCR amplification step was then used to add the index adapter sequences to create the final cDNA library. The adapter indices enabled the multiplexing of the libraries, which were pooled prior to cluster generation using a cBot instrument. The loaded flow-cell was then paired-end sequenced (76 + 76 cycles, plus indices) on an Illumina HiSeq4000 instrument. Finally, the output data was demultiplexed and BCL-to-Fastq conversion performed using Illumina’s bcl2fastq software, version 2.20.0.422.

Unmapped paired-end sequences were tested by FastQC version 0.11.3 (http://www.bioinformatics.babraham.ac.uk/projects/fastqc/). Sequence adapters were removed, and reads were quality trimmed using Trimmomatic_0.39 (Bolger et al., 2014). The reads were mapped against the reference human genome (hg38) and counts per gene were calculated using annotation from GENCODE 31 (http://www.gencodegenes.org/) using STAR_2.7.7a (Dobin et al., 2013). Normalisation, Principal Components Analysis, and differential expression was calculated with DESeq2_1.36.0 (Love et al., 2014).

### *In vitro* characterisation of cell proteomes in response to peptides 1, 7 and 1+7 in combination

In parallel with the transcriptome analysis, cells were treated for a total 7 days prior to protein extraction. Media and treatments were replenished every other day, with the media being replaced by serum-free media for the final two days of treatment prior to protein extrication and analysis. Serum-free media containing the secretome and detached matrix was first collected, then cells were briefly washed with PBS, followed by incubation with 20 mM EDTA for 1hr at 37°C to dissociate cells from the plate-attached matrix without cleavage. A protease inhibitor cocktail (P83401-1ML, Sigma), and phosphatase inhibitor cocktail 3 (P0044-1ML, Sigma) were added to the secretome and detached matrix mixture, and this sample was kept at 4°C throughout. The cell suspension was then collected, and protease and phosphatase inhibitors added as above. This suspension was centrifuged at 5000 RCF for five minutes. The detached matrix-containing supernatant was added to the secretome mixture, while the cell pellet was snap-frozen in liquid nitrogen. PBS and protease and phosphatase inhibitors were added to the plate-attached matrix, and this was stored at 4°C. Meanwhile, the secretome and detached matrix were dialysed using Slide-A-Lyzer MINI Dialysis Devices (88403, ThermoFisher Scientific) according to manufacturer’s instructions. This was then frozen at -80°C for 30 minutes, then freeze-dried overnight.

An 8 M urea buffer, containing 25 mM DTT to reduce samples, was used to resuspend the freeze-dried detached matrix and secretome. The same buffer was used separately to resuspend the cell pellet. These samples were then ultrasonicated using the Covaris LE220-plus (500 Watts peak power, 20% duty factor, 200 cycles per burst, 180 second duration). Following ultrasonication, PBS was removed from the plate-attached matrix, and the detached matrix and secretome were added to these wells, forming the total ECM proteome. All samples were brought to 10 mM iodoacetamide and kept at room temperature in the dark for 30 minutes to alkylate samples. Urea in the sample was diluted to 2 M by addition of a solution containing 25 mM ammonium bicarbonate and 1.3 mM calcium chloride. Trypsin SMART Digest Beads (60109-101, ThermoFisher Scientific) were added to all samples, then incubated overnight at 37°C to digest samples.

Trypsin SMART Digest Beads were removed using TELOS filtration tips (900-0010-096MP, Kinesis). Samples were acidified using 5 μl of 10% formic acid. Samples were mixed vigorously with ethyl acetate, to remove surfactants, glycation products, and contamination from polyethylene glycol and plastic products, then centrifuged. The upper layer was removed, and this step was repeated. The resulting aqueous bottom phase was then vacuum dried to a minimal volume. Injection solution (5% acetonitrile in 0.1% formic acid) was added to make the peptides up to 200 μl. The concentration of peptides was measured by a Direct Detect Spectrophotometer (Merck, Darmstadt, Germany), and solutions were made up to 100 μg in 100 μl of injection solution prior to desalting.

OLIGO R3 beads (1-339-03, Applied Biosystems) were mixed with an equal volume of 50% acetonitrile, and were added to a 96-well filter plate with 0.2 μm polyvinylidene difluoride (PVDF) membrane (3504, Corning). These were washed, first with more 50% acetonitrile, then with 0.1% formic acid (wash solution). The digested protein samples were then added, followed by washing twice with the wash solution. Samples were then eluted with a solution of 50% acetonitrile and 0.1% formic acid. Desalted samples were then vacuum-dried to minimal solution, and submitted to the Biological Mass Spectrometry Core Facility, Faculty of Biology, Medicine and Health, University of Manchester.

### Proteomic data analysis

LC-MS/MS was used to analyse all peptide treatment samples. This was performed by the Biological Mass Spectrometry Core Facility, Faculty of Biology, Medicine and Health, University of Manchester according to their protocols. Samples were run through the ThermoFisher Scientific Orbitrap Elite Mass Spectrometer, or through the ThermoFisher Scientific Q Exactive HF Mass Spectrometer. Following LC-MS/MS, peptide and protein identification was performed using either Progenesis QI, or Proteome Discoverer.

Progenesis QI (Nonlinear Dynamics, Waters, Newcastle, UK) was used to relatively quantify protein abundance for the initial proteomic screens of eight candidate peptides (Fig. 3) and the first proteomic analysis of P1, P7 and P1+P7 (Fig. 4c and d). Raw mass spectra files were imported and ion intensity maps were generated. Ion outlines were automatically aligned to a single reference run using default settings. Ion peaks and their relative abundances were then automatically picked without filtering and normalised to a single reference run using default settings. Data were then exported and searched using Mascot v2.5.1. Mascot was set up to search the Swiss-Prot database (selected for Homo sapiens, version 2022-08-03, 207,304 entries) assuming the digestion enzyme was trypsin. Mascot was searched with a fragment ion mass tolerance of 0.02 Da and a parent ion tolerance of 10.0 ppm. The Mascot search included peptides of charges 2+ and 3+, and a maximum of one missed cleavage site. Carbamidomethyl of cysteine was specified in Mascot as a fixed modification, while oxidation of methionine, lysine, proline and arginine were specified in Mascot as variable modifications. This was then re-imported back into Progenesis QI, where identified peptide ions were matched. Normalised abundance for each protein was calculated as the sum of each matched peptide ion abundance. Normalised protein abundances, compared between treatments, were statistically analysed within Progenesis QI using a paired (repeated measured) ANOVA test.

Relative quantification of protein abundance for the final *in vitro* proteome study (Fig. 4g) was performed using Proteome Discoverer (version 2.5.0.400; ThermoFisher Scientific, Waltham, Massachusetts, United States). Raw mass spectra files were imported, and underwent an initial processing step, using Sequest HT to identify peptides within the spectra, searching the Swiss-Prot database (selected for Homo sapiens, version 2022-08-03, 207,304 entries), and assuming the digestion enzyme was trypsin. The Sequest HT search was performed with a fragment mass tolerance of 0.02 Da, and a precursor mass tolerance of 10 ppm. Carbamidomethyl of cysteine was specified as a fixed modification. Oxidation of methionine, lysine, proline and arginine were specified as variable modifications. Following the processing step, peptide spectrum matches (PSMs) were assigned confidences based on target false discovery rates (with confidence thresholds for false discover rates set to 0.01 (strict) and 0.05 (relaxed), and adjusted p values were calculated. In parallel, protein abundances were calculated from ion intensities, and normalised abundances between treatments were compared using a pairwise ratio based t-test.

### Toxicology assessments

Prior to testing for *in vivo* effects in humans the peptide blend was solubilised into an excipient for use in topical formulations (water, pentylene glycol and propanediol). The peptides (at a concentration of 500ppm) in this excipient were then assessed in a range of *in vitro* and *ex vivo* toxicology assessments required for novel cosmetic compounds, following OECD guidelines (Development). These included a direct Peptide Reactivity Assay (DPRA) and KeratinoSens ™ (Andreas et al., 2011) to assess potential skin sensitisation (OECD No. 442C and 442D respectively), SkinEthic ™ (Alépée et al., 2010) to determine the potential skin irritation (OECD No. 439) and EpiOcular ™ (Kaluzhny et al., 2011) to assess potential eye irritation (OECD No.492).

### *In vivo* patch test of peptide combination

The peptide formulation was constituted by dissolving 1% w/w of a 50:50 mixture of P1 (500ppm) and P7 (500ppm) in a suitable vehicle to give final total peptide concentrations of either 10ppm or 30ppm. The vehicle was composed of solvent excipients (water, pentylene glycol and propanediol) added to a simple oil in water cosmetic emulsion containing water, glycerine and thickened with ammonium acryloyldimethyltaurate /VP copolymer, xanthan gum and preserved with phenoxyethanol). A vehicle control was also formulated with identical composition, including the excipients but without the peptides. Eight healthy, white but photoaged volunteers with visible signs of photoageing were recruited (Fitzpatrick skin phototype I-III; 3 male, 5 female, age range: 72-84 years old) and subjected to an extended 12-day patch test as described by Watson et al. 2008 (Watson et al., 2008). The assay uses occlusion to carry actives into the skin, so mimicking long-term use (Watson et al., 2008). The peptide formulation was applied for twelve-days, with reapplication every 4-days, alongside a vehicle control and an occluded but untreated area (baseline). The vehicle and peptide formulations (30 μl) were applied separately to the extensor photoaged forearm under standard 6mm diameter Finn Chambers (Epitest Ltd). All-*trans* retinoic acid (0.025%; Retin-A) was applied to skin for the final 4-days only (to avoid potential complications of irritation from extended occlusion) as a positive control.

On day 12, the Finn chambers were removed and 3mm punch biopsies obtained under 1% lignocaine local anaesthesia from each test site. Biopsies were embedded in optimal cutting temperature compound (Miles Laboratories, Elkhart, IN, USA), snap frozen in liquid nitrogen and stored at –80°C, prior to cutting at 10μm thickness. The study was conducted in accordance with the principles of the Declaration of Helsinki, with written informed consent (Manchester University Research Ethics Committee reference: 2020-7062-13677). Immunofluorescence analysis was performed to assess fibrillin-rich microfibril abundance (a dermal repair marker).

Fibrillin-rich microfibril abundance was quantified from sections fixed with 4% paraformaldehyde (PFA) for 10 minutes, hydrated with Tris-buffered saline (TBS; 100[1mM Tris, 150[1mM NaCl) and blocked with primary antibody diluent (Life Technologies) for 30 minutes at room temperature. Monoclonal mouse anti-human fibrillin-rich microfibril (clone 11C1.3) antibody (MA5-12770; Invitrogen) was applied overnight at 4°C. The sections were washed with TBS and incubated with VectaFluor™ Excel Amplified Anti-mouse IgG Dylight™ 488 secondary antibody (DK-2488, Vector Laboratories) for 30 minutes at room temperature. Microfibrils were visualised with the Olympus pE-300 microscope. Using ImageJ software, an area 30μm from the DEJ into the papillary dermis was analysed by thresholding the image and measuring the percentage coverage of positive fibrillin-rich microfibril staining. Differences between the test conditions were compared and analysed by repeated measures analysis of variance (RM one-way ANOVA) with Dunnett’s test where significance was p < 0.05 with GraphPad Prism software (version 9.3.1).

For extraction of biopsy RNA, once obtained, biopsies were immediately stored in RNAprotect Tissue Reagent (76104, Qiagen) to stabilise RNA. Biopsies were incubated at 37°C for one hour with 20mM EDTA, then the epidermis was separated from the dermis using forceps. Epidermal and dermal samples were then processed for RNA extraction using the RNeasy Fibrous Tissue Mini Kit (74704, Qiagen) according to manufacturer’s instructions. Briefly, samples were removed from the RNAprotect Tissue Reagent, and added to appropriate volumes of buffer RLT and homogenised at 50 Hz for 5 minutes using the TissueLyser LT (Qiagen), until all tissue was homogenised. Samples were then incubated with Proteinase K at 55°C for 10 minutes, prior to centrifugation to remove debris. A half-volume of ethanol was added, then samples were transferred to spin columns, where they were washed with buffer RW1. DNase I was then added to spin columns, and incubated at room temperature for 15 minutes. Samples underwent three further wash steps, once with buffer RW1, and the final two with buffer RPE. RNAse-free water was then used to elute RNA samples. Epidermal and dermal RNA samples were then combined and further processed to concentrate RNA and remove protein contaminants using RNeasy Mini Kit as above. Total RNA was then submitted to the Genomic Technologies Core Facility, Faculty of Biology, Medicine and Health, University of Manchester, and processed and analysed as previously.

## Acknowledgments

This work was supported by a programme grant from No7 Beauty Company, Walgreens Boots Alliance to MJS, and REBW. We thank Leo Zeef and Andy Hayes of the Bioinformatics and Genomic Technologies Core Facilities at the University of Manchester for providing support with regard to RNA-Seq. We would also like to thank the Biological Mass Spectrometry Core Research Facility in the Faculty of Biology, Medicine, and Health at The University of Manchester (RRID: <u>SCR_020987</u>).

## Author contributions

M.J.S., M.B. and EB conceptualized and managed this study. M.O. developed the bioinformatics pipeline. N.J., M.O., A.E., B.M., Y.D., C.R., L.B. and A.P., carried out the experiments. N.J., M.O., A.E., B.M., L.Z., E.B., Y.D., C.C., R.L., O.P., P.M. and M.J.S analysed the data. N.J., M.J.S., M.O., A.E., E.B., B.Z., R.E.B.W. and M.B. drafted and edited the manuscript. L.D., R.L., O.P., P.M. and C.C. edited the manuscript.

## Additional information

The identity of proteins significantly up- or down-regulated by peptides P1 and P7 in the initial in vitro proteomics screen are reported in Supplementary Item 1. The GO-term processes for all transcriptomics screens are reported in Supplementary Item 2.

Proteomics data are available via ProteomeXchange.

“Prediction, screening and characterisation of novel bioactive, tetra-peptide matrikines - 8 peptides”, Accession: PXD039941, DOI: 10.6019/PXD039941, Reviewer account: Username, Password: yhg6WW3T;

“Prediction, screening and characterisation of novel bioactive, tetra-peptide matrikines - 2 peptides and combo” Accession: PXD039929, DOI: 10.6019/PXD039929, Reviewer account: Username: , Password: tRg1OMZT.

“Prediction, screening and characterisation of novel bioactive, tetra-peptide matrikines - peptide combination, n=3”, “Accession: PXD039960, DOI: 10.6019/PXD039960, Reviewer account: Username, Password: 6FhWHe5r.

The three transcriptomics data sets (E-MTAB-12711, E-MTAB-12710 and E-MTAB-12704) can be found at ArrayExpress (https://www.ebi.ac.uk/biostudies/arrayexpress).

The peptide prediction code can be found at: https://github.com/maxozo/Matrikine_Discovery.

