## Supplemental figures and tables for "Novel in-silico predicted matrikines are differential mediators of in vitro and in vivo cellular metabolism"

| Gene | Protein | Gene | Protein |
|---|---|---|---|
| Susceptibility targets | | | |
| Protease susceptible (predicted cleavage sites) | | UVR/ROS susceptible (amino acid composition) | |
| ADM2 | Protein ADM2 | DMBT1 | Deleted in malignant brain tumors 1 protein |
| APOA4 | Apolipoprotein A-IV | DPT | Dermatopontin |
| BGN | Biglycan | EFEMP2 | EGF-containing fibulin-like extracellular matrix protein 2 |
| COLQ | Acetylcholinesterase collagenic tail peptide | EMCN | Endomucin |
| COL15A1 | Collagen alpha-1(XV) chain | FBN1 | Fibrillin-1 |
| COL17A1 | Collagen alpha-1(XVII) chain | FBN2 | Fibrillin-2 |
| CTHRC1 | Collagen triple helix repeat-containing protein 1 | FBLN1 | Fibulin-1 |
| CRH | Corticoliberin | FBLN5 | Fibulin-5 |
| FGG | Fibrinogen gamma chain | FGG | Fibrinogen gamma chain |
| IGKV2D-28 | Immunoglobulin kappa variable 2D-28 | GPX3 | Glutathione peroxidase 3 |
| LRG1 | Leucine-rich alpha-2-glycoprotein | HSPA5 | Endoplasmic reticulum chaperone BiP |
| PRELP | Prolargin | HSPD1 | 60 kDa heat shock protein, mitochondrial |
| SDC4 | Syndecan-4 | KRTDAP | Keratinocyte differentiation-associated protein |
| SERPINA6 | Corticosteroid-binding globulin | LAD1 | Ladinin-1 |
| SERPINF1 | Pigment epithelium-derived factor | MATN2 | Matrilin-2 |
| SPX | Spexin | SCGB1D2 | Secretoglobin family 1D member 2 |
| TGFB1 | Transforming growth factor beta-1 proprotein | SPX | Spexin |
| TNFSF10 | Tumor necrosis factor ligand superfamily member 10 | SRPX | Sushi repeat-containing protein SRPX |
| SRGN | Serglycin | TMSB4X | Thymosin beta-4 |
| VWA5B1 | von Willebrand factor A domain-containing protein 5B1 | UCN | Urocortin |
| ECM targets | | | |
| Collagens | | | |
| COL1A1<br>COL1A2 | Collagen I (alpha-1 and alpha-2 chains) | COL4A1<br>COL4A2<br>COL4A3<br>COL4A4<br>COL4A5<br>COL4A6 | Collagen IV (alpha-1 to alpha 6 chains) |
| COL3A1 | Collagen III (alpha-1 chain) | | |
| COL6A1<br>COL6A2<br>COL6A3 | Collagen VI (alpha-1 to alpha 3 chains) | | |
| COL7A1 | Collagen VII (alpha-1 chain) | | |
| Elastic fibre associated | | | |
| ELN | Elastin | MFAP2 | Microfibrillar-associated protein 2 |
| EMILIN1 | EMILIN-1 | MFAP4 | Microfibril-associated glycoprotein 4 |
| FBN1 | Fibrillin-1 | LTBP2 | Latent-transforming growth factor beta-binding protein 2 |
| FBLN5 | Fibulin-5 | LTBP4 | Latent-transforming growth factor beta-binding protein 4 |
| Adhesive glycoproteins and proteoglycans | | | |
| DCN | Decorin | LAMA2 | Laminin subunit alpha-2 |
| FMOD | Fibromodulin | LAMA3 | Laminin subunit alpha-3 |
| FN1 | Fibronectin | LAMA4 | Laminin subunit alpha-4 |
| LAMB1 | Laminin subunit beta-1 | LAMA5 | Laminin subunit alpha-5 |
| LAMB2 | Laminin subunit beta-2 | LAMC1 | Laminin subunit gamma-1 |
| LAMB3 | Laminin subunit beta-3 | LAMC2 | Laminin subunit gamma-2 |

**Supplemental information Table S1. Putative targets for matrikine generation in the human skin proteome**. An initial cohort of 73 proteins with the potential to be sources of matrikines *in vivo* was defined by identifying, within the cohort of extracellular human skin proteins: 20 proteins with the highest proportion of predicted cleavage sites/ protein length) i.e. the most protease susceptible and 20 proteins with the highest proportion of UVR/ROS susceptible amino acid residues. These list of susceptible proteins was supplemented with additional skin ECM targets including five structural collagens (comprising 13 alpha chains), eight elastic fibre associated proteins and twelve adhesive glycoproteins / proteoglycans. Four proteins (indicated in red: FGG, SPX, FBN1 and FBLN1) were found in more than one category leading to a final cohort of 69 target proteins.

| Pep. | Sequence | Source proteins containing 100% homologues (potentially cleaved) | Source proteins containing 100% homologues (potentially uncleaved) |
|---|---|---|---|
| **P1** | GPKG | ADIPOQ; COL1A1 (x4); COL2A1 (x10); COL3A1; COL4A1 (x6); COL4A2 (x6); COL4A3 (x2); COL4A4 (x5); COL4A5 (x3); COL4A6 (x10); COL5A1 (x2); COL5A2 (x7); COL6A2 (x7); COL6A5; COL14A1 (x2); COL15A1 ; COL16A1 (x11); COL17A1 (x5); COL18A1 (x3); COLQ (x3); CRIP2 (x3); FLNA (x4) [22] | C1QB; C1QC; CCT2; COL1A2; COL6A1; COL6A3; COL6A6; COL7A1; CCT6A; FLNB; HNRNPU; HSPD1 [12] |
| **P2** | GPSG | TNC (x8); COL7A1; COL1A1; COL1A2 (x2); COL2A1 (x3) COL5A1; COL4A1; COL4A6 (x7); COL14A1 (x4); COL16A1; COL6A2; DYNC1H1 (x2); MELTF [13] | ALCAM; COL3A1; COL4A2; COL4A4; COL4A5; COL4A3; COL5A2; COL6A1; COL12A1; COL17A1; COLQ; FLNA; GAPDH; LAMA3; LAMB1; LTBP2; MUC4 [16] |
| **P3** | LSPG | A2ML1 (x2); EMILIN1; FN1 (x3); LAMB1 (x2); CD34; HPSE; RNPEP (x2); IGHG1; IGHG2; IGHG3 [10] | C4A; C4B; COPA; ECM1; FLNA; GHR; IGKV3-20; LEP; LOX; LOXL1; PLEC; SHH; THBS4; VCAN; WNT5A [15] |
| **P4** | EKGD | COL4A1; COL4A3; COL4A5; COL6A2 (x2); COL7A1 (x2); COL15A1 (x2); COL16A1 (x2); COLQ (x4); VWA5B1; TF [10] | ADIPOQ; C1QB; COL4A2; COL4A4; COL4A6; COL5A1; COL6A5; COL6A6; COL12A1; COL17A1; COL18A1; COL14A1; TNF; GBP1 [14] |
| **P5** | QTAV | FN1 (x2); FGFR3 (x3) [2] | ANGPT2; PSAP [2] |
| **P6** | LSPD | AEBP1 (x2); CFH (x6); COL6A3 (x2); COLQ (x3); CST6 (x2); FGFR4 (x2); IFI30 (x2); MMP11 (x2); VCAN (x3) [9] | ACE2; C5; CCL27; CCT6A; COL12A1; HGFAC; TLR9 [7] |
| **P7** | LSVD | C4A (x2); C4B (x2 ); FBLN1; FLNB (x2); LAMC2; LBP (x2); NID1 [7] | C4BPA; CLU; CSPG4; ITIH5; TUBA4A [5] |
| **P8** | ELED | BGN (x3); ASPN; NID1; MYH9 [4] | COL6A6; FGA; FGG; HSPG2; LAMA2( ); NECTIN1; PLEC; SOGA1 [8] |

**Supplemental information Table S2. Peptide amino acid sequence and potential skin protein source.** For each peptide the sequence is accompanied by a list of proteins from which that peptide is predicted to be cleaved and another list of proteins where flanking cleavage sites were not predicted. Some peptides were present in multiple copies in each protein (number indicated in parentheses). For example, P1 was the most ubiquitously distributed peptide; present in 44 skin proteins (22 flanked by putative cleavage sites [potentially cleaved] with some proteins containing multiple copies (i.e. 4 copies in COL1A1).

| Peptide | Sequence | Maximal non-toxic concentration (culture medium with 0% FBS) | Maximal non-toxic concentration (culture medium with 10% FBS) | Tested Concentrations Culture medium with 0% FBS | Tested Concentrations Culture medium with 3% FBS |
|---|---|---|---|---|---|
| P1 | Pal-GPKG | 5ppm | 12,5ppm | 1, 3, 5ppm | 4, 8, 12.5ppm |
| P2 | Pal-GPSG | 4ppm | 12,5ppm | 1, 2, 3, 4ppm | 4, 8, 12.5ppm |
| P3 | Pal-LSPG | 3ppm | 12,5ppm | 1, 2, 3ppm | 4, 8, 12.5ppm |
| P4 | Pal-EKGD | 12,5ppm | 12,5ppm | 1, 5, 10, 12.5ppm | 4, 8, 12.5ppm |
| P5 | Pal-QTAV | 12,5ppm | 12,5ppm | 1, 5, 10, 12.5ppm | 4, 8, 12.5ppm |
| P6 | Pal-LSPD | 7ppm | 12,5ppm | 1, 3, 5, 7pmm | 4, 8, 12.5ppm |
| P7 | Pal-LSVD | 10ppm | 15ppm | 1, 5, 7, 10ppm | 5, 10, 15ppm |
| P8 | Pal-ELED | 7ppm | 12,5ppm | 1, 3, 5, 7ppm | 4, 8, 12.5ppm |

**Supplemental information Table S3: Toxicity testing.** *In vitro* toxicity was assessed for each peptide (at 3 – 12.5ppm) using primary human dermal fibroblasts (HDFs) cultured for 72 hours at 37°C with and without 10% fetal bovine serum supplementation. Cell toxicity was assessed using Hoechst 33258 solution to determine the maximum peptide concentration at which no cell toxicity was observed.

| Treatment | Procollagen-I | | Fibronectin | | Decorin | | Collagen-IV | | Hyaluronan | |
|---|---|---|---|---|---|---|---|---|---|---|
| | Exp1 | Exp 2 | Exp 1 | Exp 2 | Exp 1 | Exp 2 | Exp 1 | Exp 2 | Exp 1 | Exp 2 |
| **TGF-β1** | 74%** | 179%** | 165%** | 342%** | -37%** | -70%** | 154%** | 301% ** | 60%** | 131%** |
| **P1** | 60%** | 62%** | 11% | 30%** | 13% | 26% | 30%** | -9% | -17%** | -16% |
| **P2** | 42%** | 77%** | 44%** | 94%** | -13% | -3% | 50%** | 10%* | -33%** | -32%** |
| **P3** | 25%** | -9% | 82%** | 3% | 39%** | -7% | 50%** | -3% | -26%** | -42%** |
| **P4** | 52%** | 93%** | 81%** | 121%** | 57%** | 43%** | 40%** | 45%** | -1% | +6% |
| **P5** | 56%** | 59%** | 38%** | 87%** | 23%* | 33%* | -9% | 0% | -1% | 0% |
| **P6** | 29%* | 82%** | 65%** | 92%** | 1% | 24%* | 7% | 43%** | -40%** | -11% |
| **P7** | 57%** | 19%** | -9% | -15% | 43%** | 169%** | 7% | -15%** | -13%* | -8% |
| **P8** | 57%** | 10% | 70%** | 53%** | 55%** | 41%** | 42%** | 9% | +28%* | -1% |

**Supplemental information Table S4: ELISA data for 8 peptides selected for further screening.** The results from the optimum concentration for each peptide is shown (strongest results overall against the 5 ECM markers tested via ELISA: (n=2 with five experimental replicates for all proteins apart from collagen IV with four experimental replicates). Human fibroblast cell cultures were exposed to peptides P1 to P8. Two independent experiments were performed and production of procollagen-I, fibronectin, decorin, collagen-IV and of hyaluronan was measured. Data are reported as percentage changes against solvent control (0.1% of DMSO) with significant differences indicated by * ($p<0.05$) and ** ($p<0.01$).

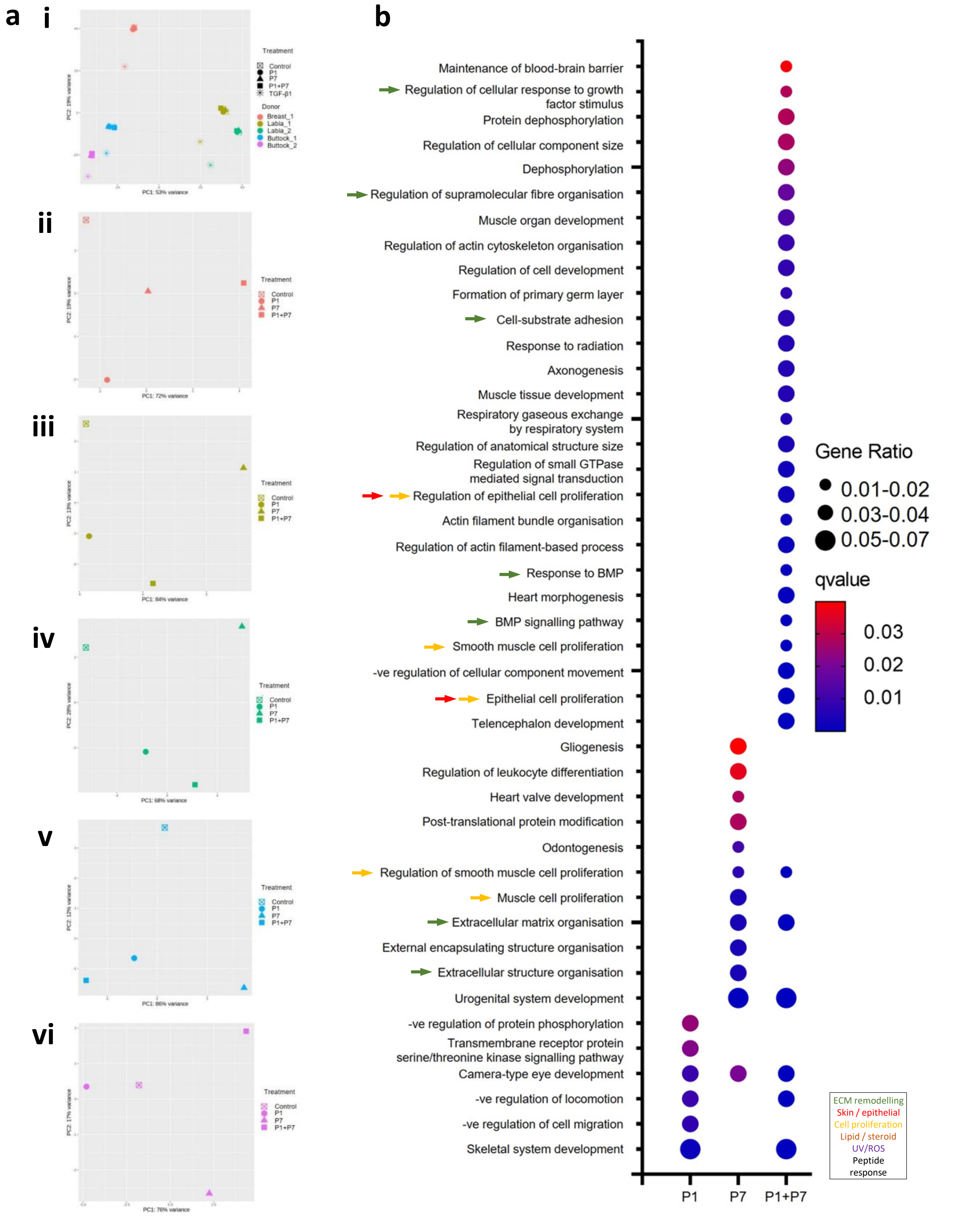


**Supplemental information Figure S1: *In vitro* characterisation of peptides 1 (PQ), 7 (P7) and combination (P1+P7).** (a) Principal component analysis (PCA) of RNA-Seq data for TGF-β1 (i) and peptide exposed HDFs derived from breast (ii), labia (ii and iii) and buttock (v and vi, n= 5 biological replicates per treatment. TGF-β1 treatment profoundly affected the transcriptomes of all donor cells compared with control. For each donor-derived cell, the peptide treatments segregated from each other and the control. (b) GO-term enrichment analysis for ECM genes of interest (27 genes defined in Extended Data Table 1: collagens, elastic fibre associated and adhesive glycoproteins and proteoglycans). Both P1 and P7 individually enriched multiple biological processes but P1+P7 in combination enriched 32 processes including those associated with ECM remodelling (green arrows) and with cell proliferation (yellow arrows).

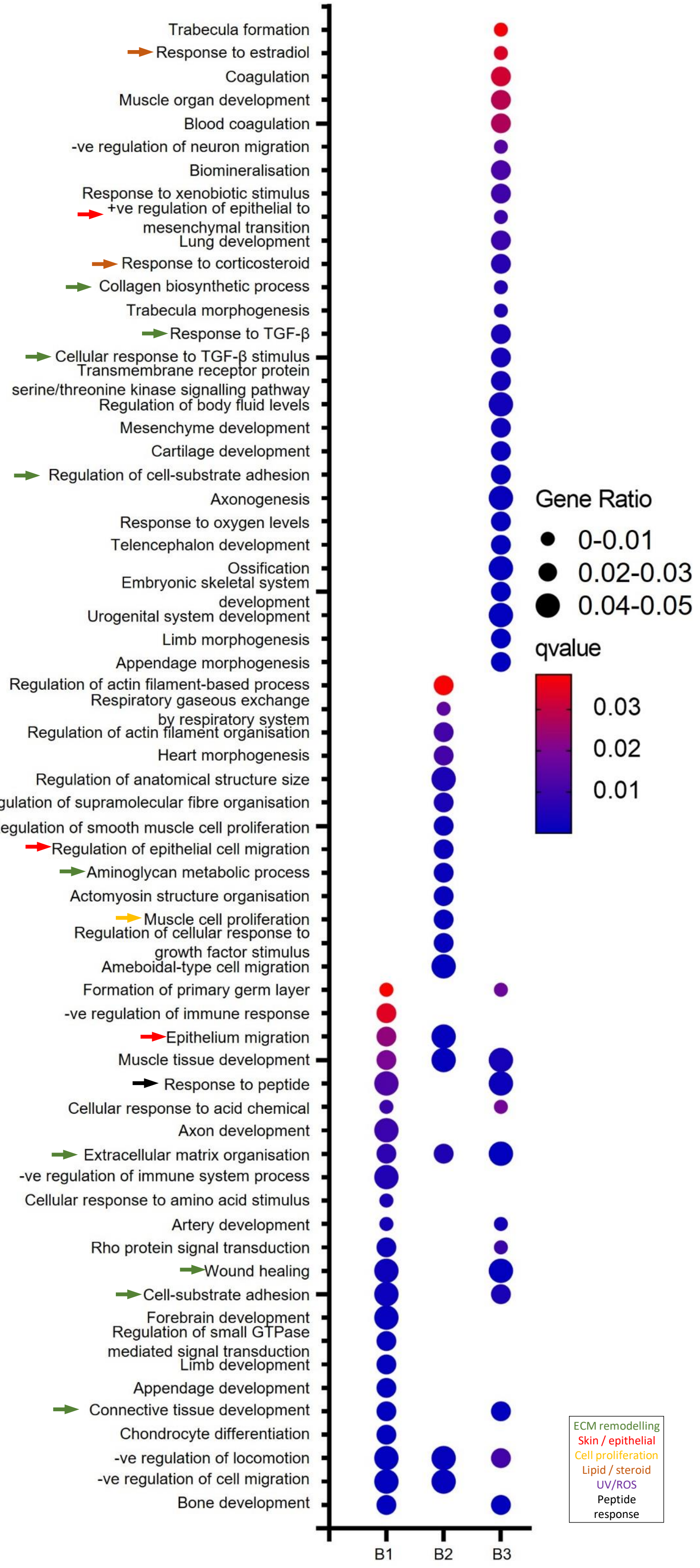


**Supplemental Figure S2: *In vitro* characterisation of peptide combination (P1+P7).** GO-term enrichment analysis for ECM genes of interest (defined in Extended Data Table 1), n= 3 biological replicates with 3 experimental replicates per treatment. The peptide combination promoted ECM organisation in HDFs derived from all three donors along with other ECM-tissue related processes in at least 2/3 donors (green arrow). For cell donors B1 and B3 the peptide combination enriched the response to peptide process (yellow arrow).
